# Inferring non-coding RNA regulatory network from transcriptomic data and curated databases

**DOI:** 10.1101/2025.04.10.648218

**Authors:** Hongjie Ke, Zhenyao Ye, Li Feng, Zhangchi Xu, Rong Pan, Eric Li, Jianfei Qi, Shuo Chen, Menglu Liang, Tianzhou Ma

**Author notes:** **Correspondence:** Tianzhou Ma, Department of Epidemiology and Biostatistics, University of Maryland School of Public Health, College Park, Maryland, 20742, U.S.A.

## Abstract

Non-coding RNAs (ncRNAs) have played an indispensable role in regulating gene expression and cellular processes. However, the complex regulatory circuits between ncRNAs and coding genes remain understudied. Major challenges for inferring ncRNA regulatory network (NRN) on a transcriptome-wide scale include high dimensionality of both ncRNA and coding gene expression data, their context-dependent interaction, and lack of validation. We hereby propose a comprehensive analytical framework, namely Construction and Analysis of non-coding RNA regulatory NETwork (*CAR-NET*), specifically designed to infer NRN from transcriptomic data and curated databases on experimentally validated ncRNA-gene interactions and disease-related ncRNAs. At its core is a novel computationally efficient Bayesian network structure learning method that leverages the semi-bipartite graph structure of NRN for dimension reduction, edge orientation and global network search. We also developed and applied algorithms to perform differential network, subnetwork and pathway analysis of the inferred network. We showed the strength of *CAR-NET* in simulations and three case studies, encompassing lncRNA regulation in brain development, miRNA differential regulation in renal cell carcinoma and regulation by different classes of ncRNAs in different cell lines using bulk or single cell RNA-seq data. An R-shiny app (https://github.com/kehongjie/CAR-NET) was developed to implement the method(s) with interactive GUI.

## Introduction

Non-coding RNAs (ncRNAs), including long non-coding RNAs (lncRNAs), micro RNAs (miRNAs), small nuclear RNAs (snRNAs), etc., are critical regulators that control the gene expression at multiple levels^1,2^. Compared to their protein-coding gene counterparts, ncRNAs are generally understudied. Revealing how the ncRNAs regulate their target genes and the complex regulatory circuits they generate provides deeper insights into the regulation mechanisms of life by ncRNAs and their roles in developmental processes and diseases^3^. A gene regulatory network (GRN) has been widely used to model the complex interactions between genes and their regulators. Current studies of GRNs, however, have focused on the interactions between transcription factors and genes^4,5^, few methods have been specifically designed to model the ncRNA-gene regulatory relationship and the associated network, leaving this indispensable component of systems biology enigmatic.

Researchers typically applied data-driven methods on transcriptomic data to infer large-scale GRNs^5-7^. Further, knowledge from existing literature or manually curated databases were used to synthesize core GRNs^5,8^. Constructing an ncRNA regulatory network (NRN) that involves ncRNA regulators and their regulated genes on a transcriptome-wide scale is challenging in a few aspects: Firstly, transcriptome-wide ncRNA and gene expression data are both of high-dimension (∼10-100k nodes each) with a huge number of potential connections (∼billions of possible edges). Direct application of most existing GRN methods is computationally infeasible. Secondly, the expression of ncRNA is highly variable and context-dependent (e.g. cell type, tissue and condition specific)^2,8^, so the regulatory relationship between ncRNAs and genes can also vary in different conditions, which is hard to identify with increasing network size. Thirdly, other than from the transcriptomic data, GRNs have been historically inferred from experimentally validated transcriptional factor activity and regulation events compiled in databases^5,9^, which has been challenging for ncRNA-gene regulatory interactions due to the scarcity of relevant databases. The growth of publicly available curated databases on ncRNA-gene interactions (experimentally determined or base-pairing prediction based) and disease-related ncRNAs (from human or animal studies) in recent years has created an unprecedented opportunity to assemble ncRNA regulatory network from both transcriptomic data and curated databases, prioritizing biologically relevant and reliable ncRNAs and ncRNA-gene interactions that will be otherwise missed due to insufficient power with transcriptomic data alone.

In this paper, we propose an analytical framework, namely Construction and Analysis of noncoding RNA regulatory NETwork (*CAR-NET*), to infer ncRNA regulatory network from transcriptomic data and curated databases on experimentally validated ncRNA-gene interactions and disease-related ncRNAs. At the core of the framework is a novel and computationally efficient two-stage Bayesian network (BN) structure learning method to infer large-scale NRN from transcriptome-wide ncRNA and gene expression data, while incorporating prior knowledge from existing curated databases. We also developed and applied algorithms to perform differential network, subnetwork and pathway analysis of the inferred network. The framework is flexible that allows input transcriptomic data from either bulk or single-cell RNA-seq (scRNA-seq) technologies, and incorporation of new curated databases on ncRNA-gene-disease interactions based on users’ interest. To the best of our knowledge, *CAR-NET* is the most comprehensive analytical framework specifically designed to infer the ncRNA-gene regulatory relationship on a transcriptome-wide scale coupled with useful subsequent biology-driven analyses to generate new hypotheses to be tested in future studies. Our tool will significantly contribute to the study of ncRNA and help biomedical researchers in ncRNA biology: 1) narrow down the transcriptome-wide ncRNA and gene expression data from bulk or single-cell RNA-seq in their study to the most critical ncRNAs and their interactions with genes at network level; 2) learn how these ncRNA-gene interactions in the network dynamically change in different conditions (e.g. clinical stages and cell types); 3) assess how well the findings from their own data align with the large amount of publicly available curated databases on experimentally validated ncRNA-gene interactions.

We showed the strength of our framework using simulations and three real data examples encompassing lncRNA regulation in brain development, miRNA differential regulation between early and late stages of renal cell carcinoma as well as the gene regulation by different types of ncRNAs in different cell lines using single cell data. To allow researchers to apply the framework easily, we also developed a user-friendly R-shiny app (https://github.com/kehongjie/CAR-NET) to implement the streamlined workflow with interactive graphical user interface (GUI).

## Results

### Overview of *CAR-NET* framework

An NRN typically has a bipartite graph backbone^10^, where one set of ncRNA nodes targets at the other set of gene nodes (Figure 1A). Direct gene-gene communications (together also known as a semi-bipartite graph) usually exist, while ncRNAs and ncRNAs are only indirectly connected via regulating the same set of gene(s), unless data for multiple types of ncRNA are available and we are specifically interested in the direct interactions between them (e.g. ceRNA network where lncRNAs interact with miRNAs^11^). Like most GRNs, NRN is sparse by nature^12^, where a majority of nodes and edges are irrelevant in the system being studied (Figure 1B). The task of inferring an NRN on a transcriptome-wide scale is to identify a few critical ncRNAs and key target genes as well as their regulatory connections in the system while taking the experimentally validated information into account.

**Figure 1.**
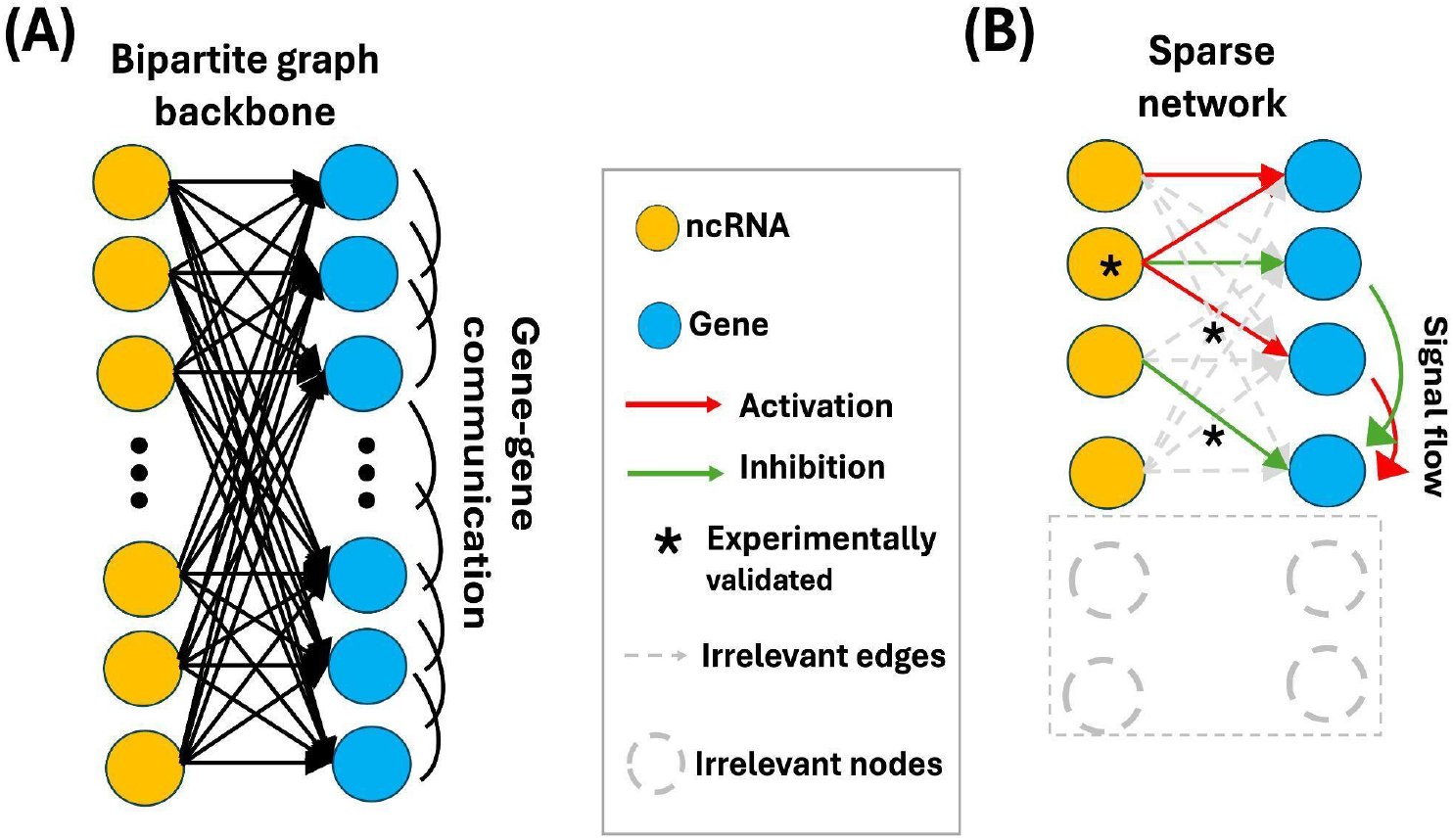
(A) A typical NRN consists of a bipartite graph backbone and gene-gene direct links (a.k.a. semi-bipartite graph). (B) The final network is sparse, including only a few critical ncRNAs and key target genes and regulatory connections. The experimentally validated information from curated database is usually taken into consideration. In other contexts, genes are used for noncoding RNAs and coding genes interchangeably. In the context of this paper, we refer genes to coding genes only.

Figure 2 shows a flow diagram of the *CAR-NET* framework. The input data include the expression data of ncRNAs and genes for matched samples (or matched cells for single cell data). The framework starts by applying standard preprocessing (e.g. normalization, transformation, filtering) to the expression data (see Methods section 4 for details). We developed a novel and computationally efficient two-stage BN learning method that leverages the semi-bipartite graph structure to construct NRN from high-dimensional ncRNA and gene expression data and curated databases (Fig 2A). In the first stage, we implemented a fast edge-wise screening that iteratively uses robust partial correlation to quickly remove noisy edges, followed by a node-wise screening that uses canonical correlation to remove nodes with no or few loosely connected edges (see Methods). After stage I, the dimensions of both ncRNAs and genes were much lower (much fewer nodes) and the network became much more sparse (much fewer edges). In the second stage, we developed a score-based order Markov chain Monte Carlo (MCMC) algorithm^13^ adapted to the semi-bipartite graph to search for the global regulatory network that best represents the data. Since the algorithm falls within a Bayesian framework, we incorporated the prior knowledge from curated databases on ncRNA-gene and ncRNA-disease relationship to prioritize disease related ncRNA nodes and reliable ncRNA-gene edges in MCMC sampling. These databases typically included experimentally validated ncRNA-gene targets^14,15^, functionally annotated^16^ or disease associated ncRNAs^17,18^(see examples in Table 1 and a complete list in Table S1). Our two-stage BN learning method will generate a final ncRNA regulatory network that best represents the data and existing curated databases.

**Table 1.**
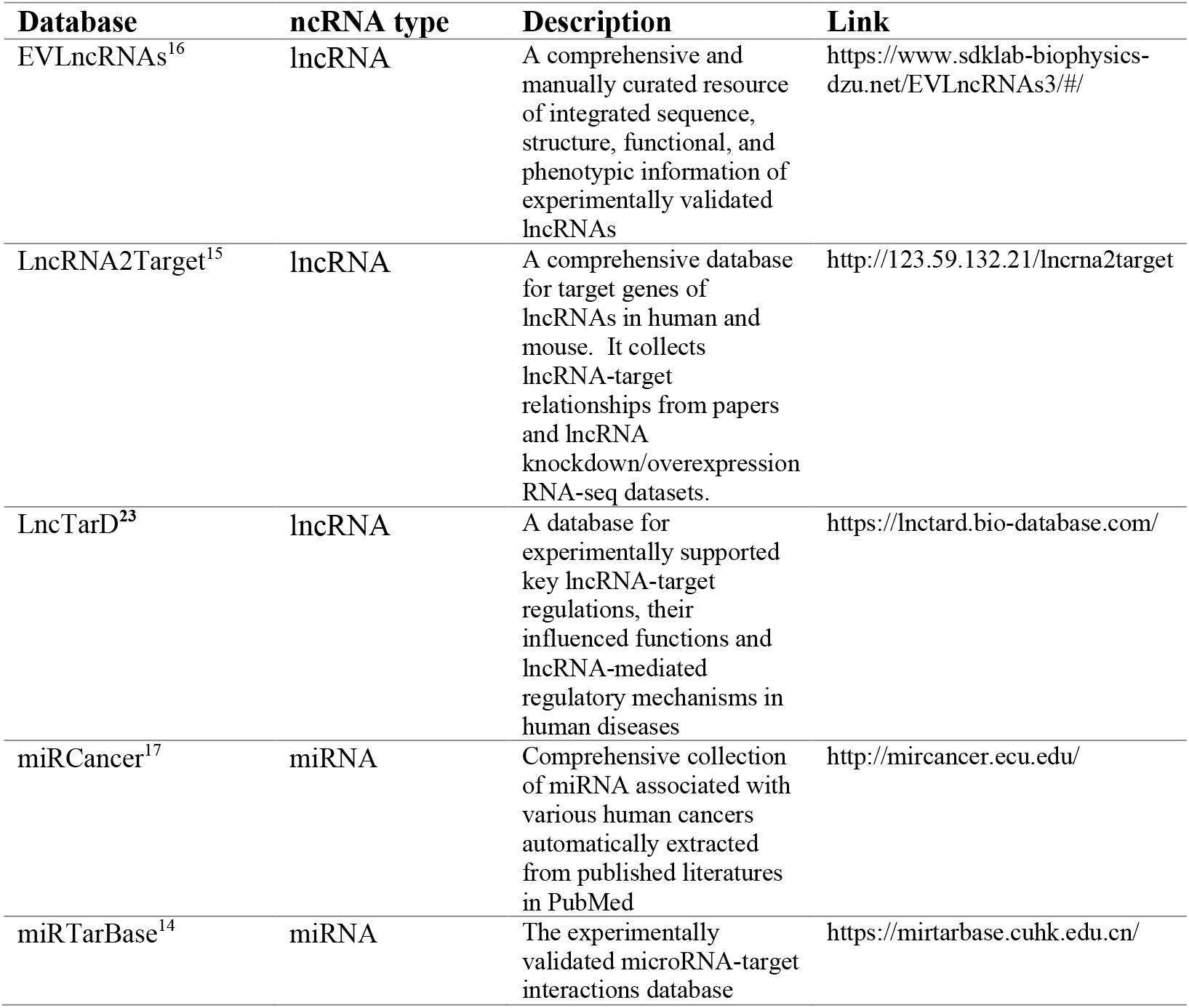
Curated databases on ncRNA-gene interactions and disease-associated ncRNAs used in case study 1 and 2.

**Figure 2.**
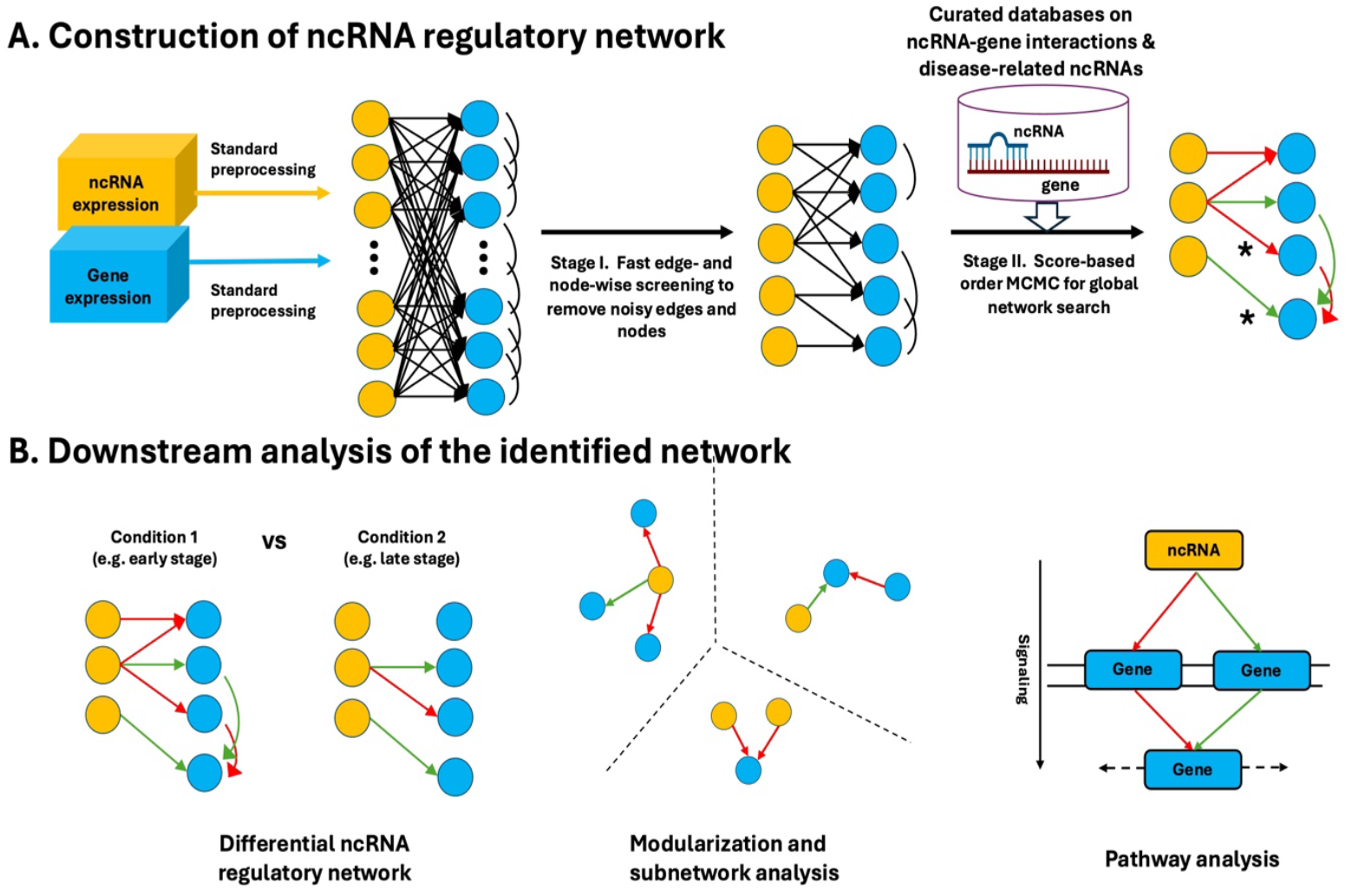
Flow diagram of *CAR-NET*. (A) A two-stage Bayesian network structure learning method to construct the ncRNA regulatory network. (B) Downstream analysis of the identified network including differential network, subnetwork and pathway analysis.

Once we identify the regulatory network, we further perform downstream analysis for more biological insights (Fig 2B). Depending on the study design, the expression data can be collected from different but comparable conditions (e.g. different clinical stages, tissues, or cell types). Detecting the differential ncRNA regulatory network can be of great interest to capture the dynamic and context-dependent interactions between ncRNA and coding genes. Direct search of a global differential regulatory network is usually computationally infeasible^19,20^ and running an algorithm independently in two conditions generates two completely different regulatory network structures that are hardly comparable. We borrowed the idea from incremental learning^21^ and proposed a computationally scalable algorithm to detect differential ncRNA regulatory network by first constructing an initial network from the reference condition and then updating to a new network using data from the other condition (see Methods). For better interpretation and more in-depth investigation of the network, we also implemented a directed Louvain algorithm^22^ to partition the identified network into smaller communities or modules for subnetwork analysis, and performed pathway analysis on the ncRNA targeted genes using curated pathway databases to highlight the biological functions targeted by these ncRNAs.

### Case Study 1: LncRNA regulation of gene expression in brain development

The important roles that lncRNAs play in brain development, neuron function, and neurodegenerative diseases are becoming increasingly evident^24,25^. Uncovering the underlying lncRNA-gene regulatory mechanism provides new insights in understanding the pathophysiology of neurodegenerative diseases with potential therapeutic values. In this example, we retrieved the lncRNA and gene expression data (bulk RNA-seq, in FPKM) of n=55 human brain samples from 4 weeks post conception to adulthood stored in LncExpDB database^26^. We followed the guideline in Sarropoulos et al^27^ to filter out lnRNAs with mean FPKM <1 and genes with mean FPKM <10 and perform log2 transformation. A total of 3,367 lncRNAs and 4,511 genes remained after preprocessing. Three external curated databases, EVLncRNAs^16^, LncRNA2Target^15^ and LncTarD^23^ (Table 1) were used to prioritize lncRNAs previously reported to regulate gene expression in the brain as well as experimentally validated lncRNA-gene interactions.

*CAR-NET* identified a final regulatory network that includes 408 lncRNAs and 588 genes (669 ncRNA->gene edges and 509 gene->gene edges, see Table S2 for a summary and Extended Data Fig 1A for the network graph). The final network we identified is more sparse than existing GRN inferring methods including PC^28^, GENIE3^29^, Pearson^30^ or WGCNA^31^ (see Methods) with fewer edges and nodes, and includes a much higher proportion of the backbone ncRNA-gene (level-1) edges than the gene-gene (level-2) edges (Fig 3A-C, Table S2). This is consistent with our simulation results (Extended Data Fig 4 and Supplementary section S2) where *CAR-NET* generated a more sparse network without losing most of the true signals. The genes being regulated by lncRNAs in the network were enriched in 28 significant pathways (BH adjusted p-value <0.05) including those specific to brain development such as endosome^32^, dendrite^33^ and myeloid cell differentiation^34^ (Extended Data Fig 1B and Table S3). We detected a total of 23 modules (>=10 nodes) from the final network using the Louvain algorithm. Out of the three largest modules detected (Extended Data Fig 1C), we identified one representative module (Fig 3D) featured by two lncRNAs belonging to the Small nucleolar RNA host gene (SNHG) class (*SNHG6, SNHG19*, also found in the experimentally validated EVLncRNA database as highlighted in *). They have been previously reported to regulate genes that encode the ribosomal protein (e.g. *RPS15, RPS27*) related to nervous system development^35^ and *BAX* gene that regulate apoptosis in the developing brain^36^. When we ordered the samples by the collection time in a time series plot, we found these two lncRNAs and the genes they regulated co-expressed with a few common spikes in 6-12 weeks post-conception, 4 days to 2 years post-birth as well as 7-14 years post-birth, all critical periods for brain development (Fig 3E-F, highlighted)^27,37^.

**Figure 3.**
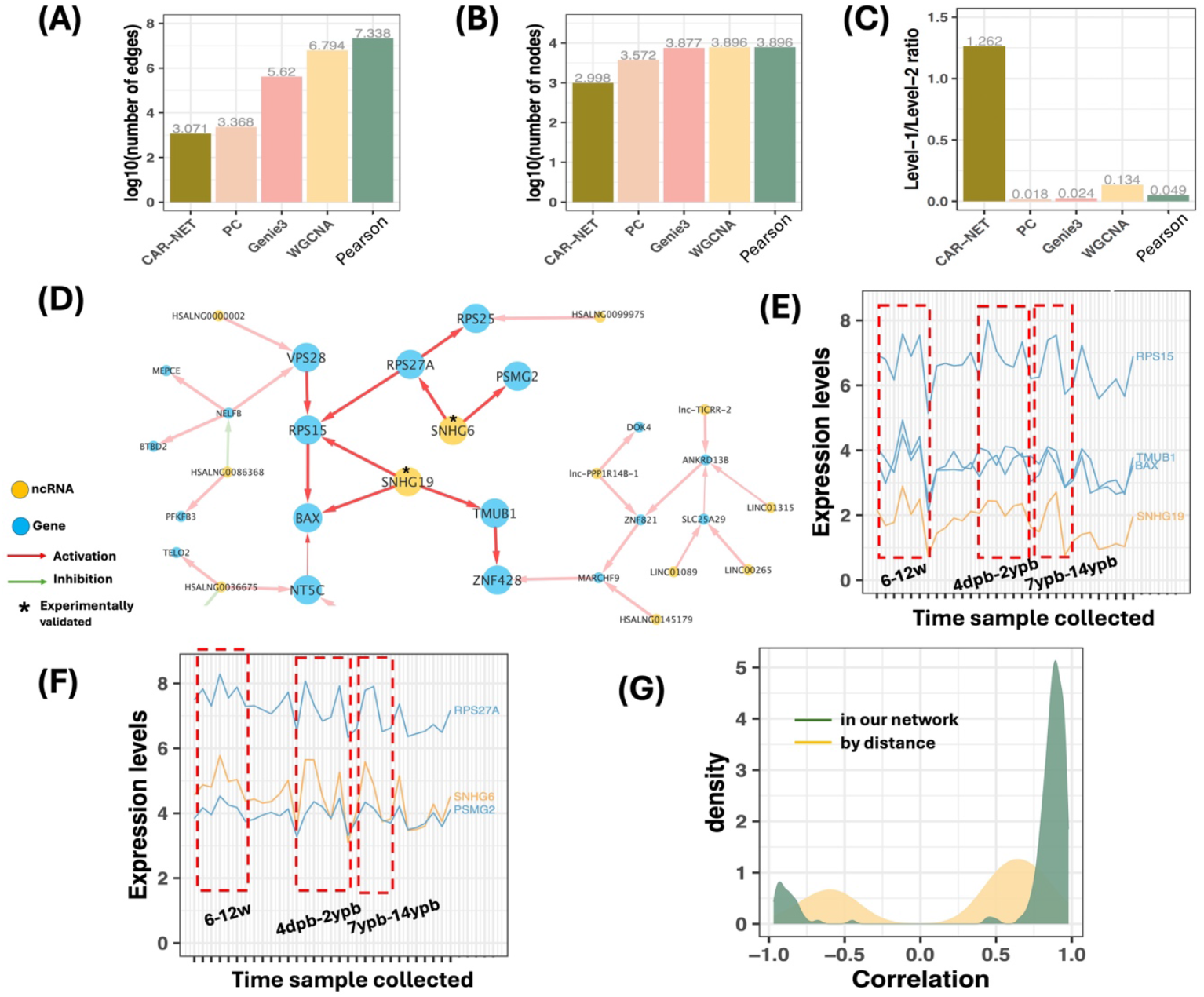
*CAR-NET* identifies a key module in lncRNA regulatory network that regulates brain development. (A) Comparison of number of edges (on log10 scale) in the final network detected by different methods. (B) Comparison of number of nodes (on log10 scale) in the final network detected by different methods. (C) Comparison of level 1 (ncRNA-gene) / level 2 (gene-gene) edge ratio in the final network detected by different methods. (D) A key module detected in our network featured by two lncRNAs in SNHG class: *SNHG6* and *SNHG19*. Line width indicates the posterior probability of the edge inferred from MCMC. “*” indicates the lncRNA nodes or lncRNA-gene edges from the curated database. (E) Time series plot ordered by sample collection time (from 4 weeks post conception to 58 years post birth) to visualize the co-expression of lncRNA *SNHG19* and its regulated genes. (F) Time series plot ordered by sample collection time to visualize the co-expression of lncRNA *SNHG6* and its regulated genes. (G) Comparison of Pearson correlations of the remaining lncRNA-gene edges in the network we identified with the edges defined by physical distance between lncRNA and gene only (i.e. cis-regulation).

It has been reported^38^ that a few well-characterized lncRNA regulated gene targets of their immediate neighbors (i.e. cis-regulation). Interestingly, in our study, in addition to the cis-lncRNA-regulators (defined as <1Mb of transcription start site; 18 out of 669), we found that many lncRNAs in our network are potentially trans-regulators (651 out of 669). The lncRNAs in the final regulatory network have much stronger marginal correlations with their regulated genes than the lncRNA regulators defined only by distance (see Fig 3G, Wilcoxon p-value=2.57e-151), implying the existence of trans-regulation of genes by distant lncRNAs and revealing the complexity of lncRNA regulation of gene expression during the organ development in human^27^.

### Case study 2: miRNA differential regulation of gene expression in early and late stages of papillary renal cell carcinoma in TCGA

Kidney cancer is among the top 20 most common cancers that represents 2-3% of all malignancies worldwide. As compared to the most common clear renal cell carcinoma, papillary renal cell carcinoma (PRCC) is heterogeneous cancer that consists of various types of renal cancer. Type 1 PRCC tumors are predominantly early stage (e.g. stage I or II), whereas type 2 tumors are frequently late stage (e.g. stage III or IV) with worse survival outcomes^39^. miRNAs play critical roles in predicting recurrence and unfavorable prognosis of renal cancer and potentially discriminate between different types of renal tumors^40^. Here, we applied our framework to study the miRNA regulation of gene expression in PRCC in TCGA Kidney Renal Papillary Cell Carcinoma (KIRP) cohort and how such regulation differs in different pathological stages. We downloaded both miRNA (in RPM) and gene expression data (in RPKM) from LinkedOmics^41^. We applied standard preprocessing by filtering out miRNAs with mean RPM <0.3 and genes with mean RPKM<5, and performing log2 transformation. The processed data include p=750 miRNAs and q=7,311 genes for n=260 matched KIRP samples. These KIRP samples are from either “early” (I-II, n=193) or “late” stage (III-IV, n=67) so we can study how the miRNA-gene regulation changes as KIRP progresses. We incorporated two curated external databases most relevant to the problem, miRCancer^17^ and miRTarBase^14^ (Table 1) to prioritize the cancer related miRNAs and experimentally validated microRNA-target gene interactions for more reliable and biologically interesting findings.

Using samples from early-stage KIRP, *CAR-NET* identified a final miRNA regulatory network that includes 127 miRNAs and 377 genes (Table S4 and Extended Data Fig 2A). The identified network is more sparse than existing GRN inferring methods with fewer edges and nodes while includes a much higher proportion of ncRNA-gene (level-1) edges (Fig 4A-C, Table S4). Genes regulated by miRNAs were enriched in typical cancer cell proliferation related pathways (e.g. amino acid metabolism, G protein-coupled receptors) and renal carcinoma specific pathways (e.g. calcium homeostasis) (Extended Data Fig 2B and Table S5). Using late-stage samples, our differential regulatory network algorithm further updated to a new network. In contrast to the network for early stage, the network for late stage has fewer edges forming isolated subnetworks (Table S4), implying the loss of key regulatory pathways by miRNAs (especially those playing as tumor suppressors) as the carcinoma progresses^42,43^. To account for the difference that stems from imbalanced sample sizes in two conditions instead of the real biological difference, we reran the algorithm by updating the network using subsamples of early stage with same sample size as late stage and only kept edges not differential in subsamples (see Methods and Extended Data Fig2C for the five largest differentially regulated modules). Fig 4D-E shows a network module with the largest difference between two stages. miR-935 and miR-598, reported in miRCancer database for their associations with multiple cancer types, were found to regulate multiple genes in early stage PRCC but lost their connections in late stage. The gene *NDN* they regulate function as tumor suppressors for a few types of cancer^44^, the gene *TMEM130* belongs to the family of transmembrane proteins critical for cell proliferation, invasion, and metastasis, and potentially serving both oncogene and tumor suppressor functions^45^. The loss of their association with miRNAs (Fig 4F-G) may imply miRNA dysregulation and downregulation of tumor suppressors that inhibit cell proliferation and tumor development as the PRCC progresses to late stage. Interestingly, these differentially regulated genes by miRNAs are not always differentially expressed between the two stages (Extended Data Fig2D-E), implying a rewiring of miRNA-gene regulation rather than simple up- or down-regulation of individual miRNAs/genes as cancer progresses.

**Figure 4.**
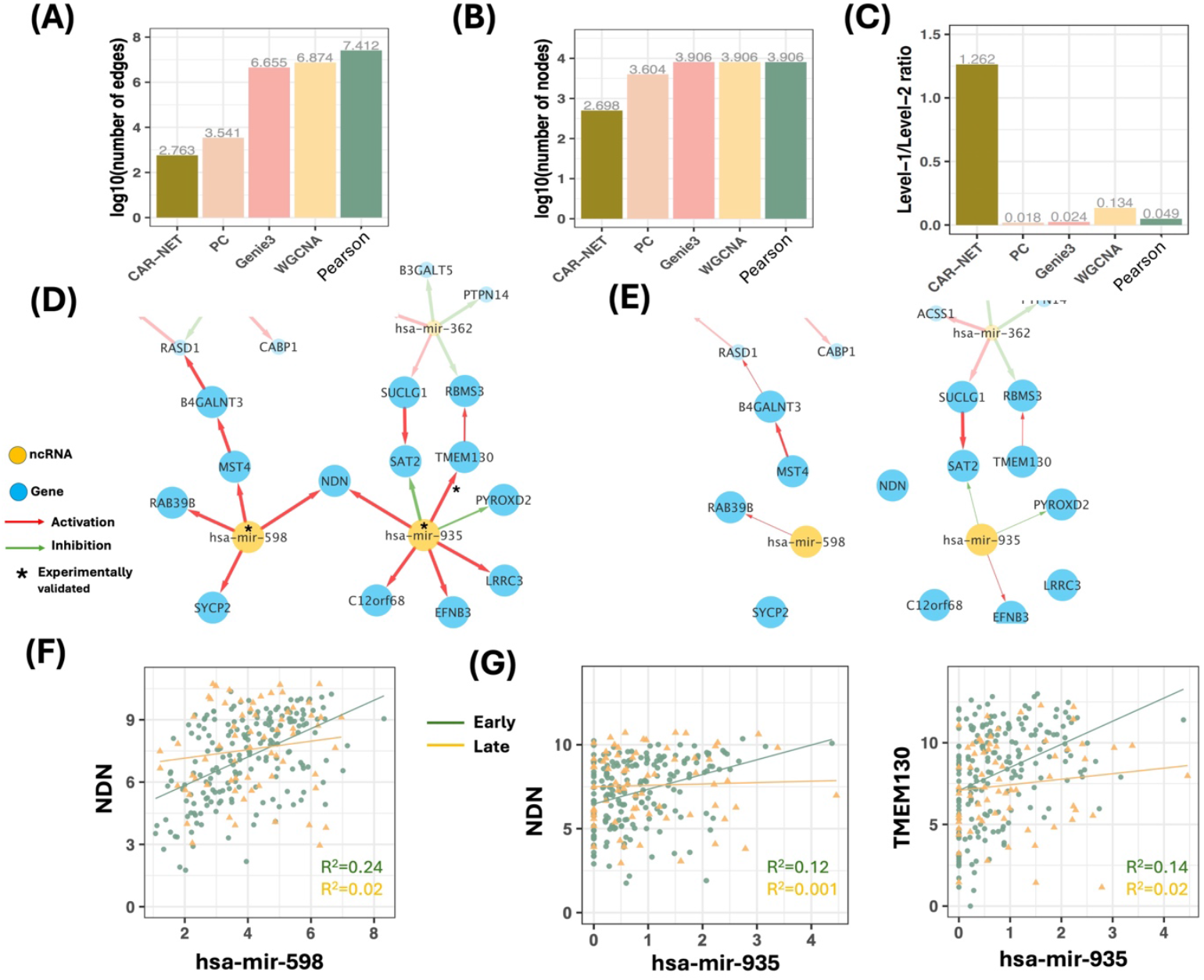
*CAR-NET* identifies differential miRNA-gene regulatory networks between early and late pathological stages of KIRP. (A) Comparison of number of edges (on log10 scale) in the final network detected by different methods. (B) Comparison of number of nodes (on log10 scale) in the final network detected by different methods. (C) Comparison of level 1 (ncRNA-gene) / level 2 (gene-gene) edge ratio in the final network detected by different methods. (D, E) Representative module inside the network that shows differential miRNA-gene regulation between early (D) and late (E) stages. Line width indicates the posterior probability of the edge inferred from MCMC. “*” indicates the lncRNA node or edge from the curated database. (F) Scatterplot of *NDN* expression vs. mir-598 expression in early and late stage of KIRP. (G) Scatterplot of *NDN* and *TMEM130* expression vs. mir-935 expression in the early and late stage of KIRP.

### Case study 3: Regulation of gene expression by diverse types of ncRNAs in different cell lines using single-cell RNA-seq data

scRNA-seq technology has become the state-of-the-art approach for unraveling the heterogeneity and complexity of RNA transcripts within individual cells^46^. A new technology called Smart-seq-total has been developed to measure a broad spectrum of coding and non-coding RNA expression simultaneously from single cells^47^. In this example, we applied our method to identify a gene regulatory network regulated by a variety of different ncRNA species (e.g. miRNA, lncRNA, snRNA and snoRNA) in different cell lines from Smart-seq-total data. We retrieved the ncRNA and gene expression data (in CPM) of three cell lines: Fibroblasts (n=278 cells), Human embryonic kidney 293 cells (HEK293) (n=260 cells) and Michigan Cancer Foundation-7 (MCF-7) (n=95 cells), from GSE151334. We filtered out ncRNA and genes with low expression and excessive zeros and performed log2 transformation following the guideline in Isakova et al. (2021)^47^ for each cell type. Due to the lack of cell line specific databases on ncRNA-gene interactions, we did not include any curated databases for this example.

We identified a multi-ncRNA regulatory network for each cell line separately (Table S6 and Extended Data Fig3A-C). Comparing across the three cell lines, ncRNA-regulated genes in fibroblasts were mainly enriched in Wnt signaling pathway critical in fibroblast differentiation, and immunity and inflammation related RIG-I and NF-κB pathways in which fibroblasts serve as key players (Extended Data Fig3D and Table S7); genes in HEK293 were enriched in metabolism related signaling pathways, fitting the cell line’s role as a model system for studying various cellular processes (Extended Data Fig3E and Table S8); genes being regulated in MCF7 were enriched in pathways related to cell cycle and its regulation due to the nature of the breast cancer cell lines (Extended Data Fig3F and Table S9). Fig 5A-C showed circular plots of edges between genes and different types of ncRNAs in the final network. lncRNA regulators were enriched in all cell lines (Table S10). For example, the lncRNA MALAT1^48^ regulated *TMSB10* for cytoskeleton organization in fibroblasts, *SRP9* involved in protein secretion and folding in HEK293, and *FTL* for iron metabolism disturbances in MCF7 (Fig 5D-F). Interestingly, some snoRNAs and snRNAs appear to be important regulators in HEK293 and MCF7 cell lines with borderline significance (Table S10 and Fig5G-H). With the overall low expression of snoRNAs and snRNAs, however, follow-up studies are needed to validate these findings. Such variation in ncRNA composition and relative importance we observed here is consistent with previous findings^47^ and has reflected the cell-to-cell heterogeneity in ncRNA regulation.

**Figure 5.**
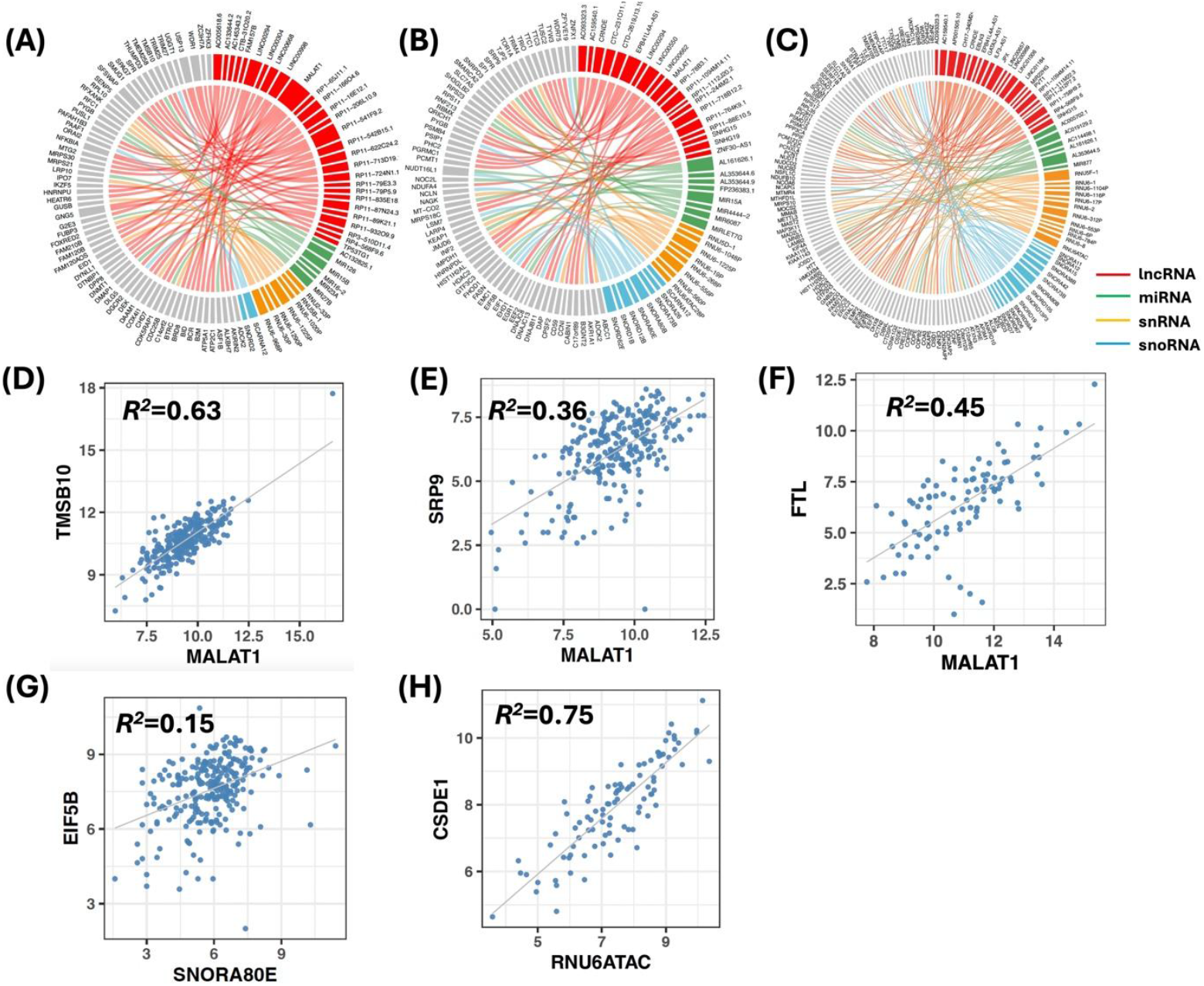
*CAR-NET* identifies the cell line specific ncRNA-gene regulatory network and reveals cell-to-cell heterogeneity. (A-C) Circular plots of genes regulated by different types of ncRNAs in fibroblasts, HEK293 and MCF7, respectively. (D) Scatterplot of selected lncRNA *MALAT1* and its regulated gene *TMSB10* in fibroblasts. (E) Scatterplot of selected lncRNA *MALAT1* and its regulated gene *SRP9* in HEK293. (F) Scatterplot of selected lncRNA *MALAT1* and its regulated gene *FTL* in MCF7. (G) Scatterplot of selected snoRNA *SNORA80E* and its regulated gene *EIF5B* in HEK293. (H) Scatterplot of selected snRNA *RNU6ATAC* and its regulated gene *CSDE1* in MCF7.

## Discussion

In this study, we proposed a novel framework called “*CAR-NET*” to infer ncRNA regulatory network on a transcriptome-wide scale from transcriptomic data as well as curated databases on ncRNA-gene-disease interactions and perform a series of biology-driven analyses on the constructed network including differential network, subnetwork and pathway analysis. To the best of our knowledge, *CAR-NET* is the most comprehensive network-based analytical framework specifically designed to model the regulatory relationship between ncRNAs and coding genes while considering the context dependent expression of ncRNAs and their dynamic regulation on genes. Compared to existing general-purpose GRN inferring methods, *CAR-NET* fully leverages the unique semi-bipartite graph structure of NRN for dimension reduction, edge orientation and global network search, and tends to identify a more sparse network and retain a higher proportion of ncRNA-gene edges. Regulatory networks inferred from expression data alone with finite sample size can be underpowered and less accurate. Moreover, ncRNAs are lowly expressed and heterogeneous, making them even more underpowered in network detection. *CAR-NET* increased the chance of sampling more reliable experimentally validated ncRNA-gene interactions from the curated databases, enhancing the accuracy and biological relevance of the inferred network.

Bipartite-type regulation is not unique nor specific to ncRNA regulation of gene expression. For example, epigenetic regulation of gene expression by DNA methylation is essentially bipartite where certain methylation sites serve as suppressors or activators of expression of their target genes. In this study, we consider the bipartite backbone and the direct gene-gene communication. ncRNA and ncRNA only interact indirectly via regulating the common genes. This assumption is intuitive when there is only one type of ncRNA available. In the presence of multiple types of ncRNAs (e.g. case study 3), for example, lncRNA can directly interact with miRNA in a ceRNA network^11^. Future development of the tool will further consider the complex interactions and cross-talks between multiple types of ncRNAs.

In this study, we have integrated the curated databases into our BN structure learning method to prioritize more reliable interactions. Incorporation of existing biological knowledge is common in the application of Bayesian methods to omics integration problems^49^. Currently, we only incorporated curated databases on experimentally validated ncRNA-gene interactions. As the AI techniques burst nowadays, more prediction-based databases on ncRNA-gene interactions, e.g. AlphaFold^50^, become available. Future development will consider incorporating these new databases, while ensuring the quality and reliability.

We also developed a novel algorithm to detect differential network by updating from an initial network based on reference condition to a new network using data from the other condition. Detecting a global differential network is NP hard problem, our new algorithm took a shortcut by updating directly from an existing reference network using new data and comparing edges on the basis of the reference network. Indeed, such an algorithm cannot guarantee finding the optimal differential network globally, but it is sufficient for practical use in identifying the local difference in network structure between two conditions. This algorithm also has the potential to be extended to compare local structure differences between more than two networks or continuously changing networks over a time course. All the methods in our framework are made available in the R-shiny app we developed (https://github.com/kehongjie/CAR-NET), with graphical and tabular outputs and interactive network visualization provided.

## Methods

### 1. Constructing ncRNA regulatory network

*CAR-NET* uses the Bayesian network (BN) model to construct an ncRNA regulatory network (NRN) from transcriptomic data of both ncRNA and coding genes, as well as the curated databases on ncRNA-gene and ncRNA-disease interactions. BN is a graphical model that represents the probabilistic relationships among a collection of variables and their conditional dependencies and has been widely used to model the gene regulatory networks^4-6^. As compared to information theory or machine learning based GRN inferring methods, methods based on BN have several strengths by directly modeling the causal relationship, quantifying the uncertainty, and naturally incorporating prior biological knowledge to infer regulatory network structure.

The BN structure *G* is a directed acyclic graph (DAG) whose nodes represent the variables and the edges represent probabilistic dependencies between them. The DAG *G* can be defined by a set of nodes *V* and a set of directed edges *E*: *G* = (*V, E*). Each node in *V* corresponds to one variable and a directed edge in *E* ⊆ *V*x*V* represents a direct conditional relationship between the two variables in the observational data. BN structure learning is computationally challenging, to date a majority of BN structure learning methods developed fall into three main categories^51,52^: constraint-based, score-based and hybrid approaches. Detailed review of the representative BN structure learning methods in each category and their pros/cons can be found in the Supplement.

In this study, we used continuous expression data as input so we will assume a Gaussian Bayesian network. Suppose the expression data of p ncRNAs and q genes for n samples are available after preprocessing, denote the ncRNA expression data as *X*_*nxp*_ = (*X*_*1*_, …, *X*_*p*_) and the gene expression data as*Y*_*nxq*_ = (*Y*_*1*_, …, *Y*_*q*_). The nodes in the DAG represent the ncRNAs and genes, and the edges represent the direct probabilistic dependencies between them. In an initial fully connected network *G*_*0*_ = (*V*_*0*_, *E*_*0*_) with semi-bipartite structure (Fig 1A), *V*_*0*_ = {*X*_*1*_, …, *X*_*p*_, *Y*_*1*_, …, *Y*_*q*_} with |*V*_*0*_| = *p* + *q*, and *E*_*0*_ = {*X*_*j*_−> *Y*_*k*_, *1* ≤ *j* ≤ *p, 1* ≤ *k* ≤ *q*}⋃{ *Y*_’_ − *Y*_’′_, *1* ≤ *l* ≠ *l*′ ≤ *q*} with |*E*_*0*_| = *pq* + *q*(*q* − *1*)/*2*, where |.| is the cardinality of a set. Our purpose is to learn the structure of the final DAG *G*_*_ = (*V*_*_, *E*_*_) for the NRN, representing the important regulatory relationship between a few critical ncRNAs and coding genes (i.e. |*V*_*_| << *p* + *q*, |*E*_*_| << *pq* + *q*(*q* − *1*)/*2*).

Motivated by the semi-bipartite graph structure of NRN (Fig 1A), we proposed a novel two-stage hybrid BN structure learning method which is both more accurate and computationally more efficient in revealing the underlying regulatory network. In the first stage (section 1.1), we performed fast constraint-based edge-wise and node-wise screening algorithms locally to remove irrelevant edges and nodes, quickly reducing the searching space of DAGs and constructing the skeleton. In the second stage, we implemented a score-based order MCMC algorithm on the skeleton (section 1.2) and incorporated the prior knowledge on ncRNA-gene and ncRNA-disease interaction information from the curated database (section 1.3) to globally search for the best regulatory network structure that represents the data and curated database.

#### 1.1 Stage I screening and skeleton construction

In the first stage, we performed fast edge-wise and node-wise screening locally to quickly reduce the number of edges and nodes (i.e. dimension reduction) while keeping the most important edges and nodes in the network^53-55^. To distinguish the two types of edges, we called ncRNA->gene (*X*_*j*_−> *Y*_*k*_) directed edges “level-1” edges and gene-gene 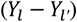 undirected edges “level-2” edges (whose direction will be inferred from our algorithm next). As the semi-bipartite graph structure is central to an NRN and level-1 edges being the backbone, we considered screening for level-1 edges with known direction first and then screening for level-2 edges, which greatly speed up the computation.

For edge-wise screening, we proposed to use our previously developed robust partial correlation (rPCor) measures^55,56^ in the context of multivariate regression (see connection in Supplementary section S1.7) as follows to perform conditional independence tests to remove irrelevant level 1 and level 2 edges, respectively:

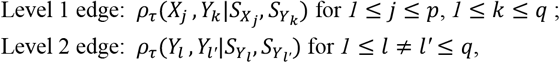

where 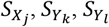 and 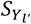 are the conditional sets of nodes (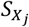include ncRNA nodes and the other three sets include gene nodes). The robust partial correlation *ρ*_τ_(*X, Y*|*S*) is defined as the correlation between the residuals resulting from the linear regression of *X* on *S* and *Y* on *S*, where *τ*is the robustification parameter that stabilizes the partial correlation estimation in the presence of heavy-tailed distribution common in transcriptomic data^57^, and is automatically selected using previously developed algorithms^58^.

Constraint-based algorithms including PC^28^ and FCI^59^ algorithms conducted conditional independence tests on partial correlation to test for the presence of edges, which can potentially remove spurious correlations. Similarly, we used a tail-robust version of partial correlation rPCor to conduct conditional independence tests to test for the presence of edges (i.e. test for 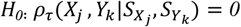 and 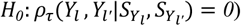 and screen out spurious correlations. Our algorithm, however, is unique in a few aspects. First, our algorithm set selection criteria for the inclusion of nodes into the conditional sets. Selection of conditional sets is critical to the accuracy and computational cost of the algorithm. Typically, we only included neighboring ncRNAs and genes marginally correlated with the nodes of interest into the conditional sets (e.g. |*ρ*| > *0*.3 according to Cohen’s guideline^60^). The same idea has been used in masked attention in Graphical Attention Network^61^. Secondly, to further improve the computation, we proposed an iterative algorithm that starts with lower order conditional independence test (e.g. when 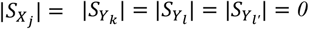, equivalent to marginal independence test) and gradually increases the order (i.e. size of conditional sets) of the conditional independence test. If any of the tests have p-values less than the threshold α, we concluded the conditional independence and screened out that edge. Low-order conditional independent tests and iterative algorithms have been shown to be efficient in high-dimensional gene expression application including the inference of gene regulatory networks^54,62^. Thirdly, unlike other constraint-based methods that used the conventional choice of α=0.05 for conditional independence test p-values without much justification. The choice of the threshold α is critical to constraint-based methods as it balances between the sensitivity and false positive control of the method.

We here proposed a fast data-driven procedure to select the optimal choice of α (Algorithm S1 in Supplementary section S1.2.2). As compared to existing constraint-based algorithms, our algorithm is much faster, less prone to error propagation and robust against heavy-tailedness. The computational complexity for edge-wise screening is of order *O*(*npq*), the same level as marginal correlation.

After screening out implausible level-1 and level-2 edges, we can identify the colliders and follow the orientation rules as in PC algorithm to orient level-2 edges in accordance with the graph structure assumed. The unique semi-bipartite graph structure of NRN enabled us to further orient the directions of some level-2 edges in addition to the routine orientation rules (see Supplementary section S1.3 and Fig S2).

The above steps only performed edge-wise screening and removed edges (assume a partially directed graph *G*. = (*V*_1_*E*_1._) left after edge-wise screening). As a by-product, nodes with all edges removed will also be screened out. However, it is common to see some nodes (either ncRNAs or genes) having very few edges connected (i.e. very small degree) were still left in the network. These nodes might be isolated (e.g. ncRNAs whose regulated genes are not linked to the core network) or at the terminal of a network (e.g. those with low betweenness centrality), thus having less influence on the full network. To guarantee more efficient computation and better convergence in the next stage, we further considered a node-wise screening by calculating the following canonical correlations between each node and its remaining edges:

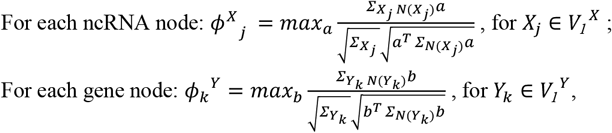

where 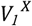 and 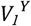 are the remaining ncRNA and gene nodes, respectively. *N*(*X*_*j*_) includes all the gene nodes linked to jth ncRNA and *N*(*Y*_*k*_) includes all the ncRNA and gene nodes linked to kth gene. 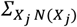 is the vector of covariance between ncRNA *X*_*j*_ and its directly linked gene nodes, and 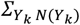 the vector of covariance between gene *Y*_*k*_ and its directly linked ncRNA and gene nodes. 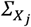 and 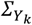 are the variances of *X*_*j*_ and 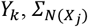 and 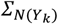 are the diagonal matrices of variances of the directly linked nodes. Nodes with small *ϕ* values will be removed, as determined by Wilks’ Lambda asymptotic test^63^. Detailed algorithm can be found in the Supplementary section S1.4. This will help remove some “scattered” nodes that are not closely connected to other nodes and won’t contribute enough to the final graph. The computational complexity of node-wise screening is of the order *O*(*np*_*1*_*q*_*1*_), where *p*_*1*_ << *p* is the number of active ncRNAs and *q*_*1*_ << *q* is the number of active genes after edge-wise screening.

#### 1.2 Stage II searching the “best” DAG

The first stage greatly reduced the number of edges and nodes thus the search space and constructed the network skeleton for the next stage to find the “best” DAG globally. In the second stage, we proposed a score-based order MCMC algorithm^13^ adapted to the semi-bipartite graph and incorporated the prior knowledge on ncRNA-disease association and ncRNA-gene interaction information to search the causal structure that best represents the data and curated database.

Suppose we had a partially directed graph *G*_*2*_ = (*V*_*2*_, *E*_*2*_) left after stage I, where 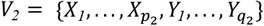, *p*_*2*_ and *q*_*2*_ are the number of remaining ncRNAs and genes, *E*_*2*_ includes all remaining edges, including both directed (ncRNA-gene directed edges and some gene-gene directed edges determined in stage I) and undirected edges (gene-gene edges not yet determined in stage I). We know that *p*_*2*_ << *p, q*_*2*_ << *q* and | *E*_*2*_| << *pq* + *q*(*q* − *1*)/*2* after stage I.

To evaluate how well a DAG represents the data, a natural strategy in Bayesian inference is to assign a score for each DAG *G* using its posterior probability given the data *D*, e.g. *P*(*G*|*D*) *∝P*(*D*|*G*)*P*(*G*), where *P*(*D*|*G*) = ∫ *P*(*D*|*G, θ* _*G*_)*P*(*θ*_*G*_ |*G*)*dθ*_*G*_ is the likelihood marginalized over the parameter space and *P*(*G*) is the graph prior and *P*(*θ*_*G*_|*G*) the parameter prior. If the graph and parameter priors (e.g. the Normal-Wishart prior) satisfy certain conditions of structure modularity, parameter independence and parameter modularity^13^, then the score (called the Bayesian Gaussian equivalence (BGe) score) for the graph *G*_*2*_ can be decomposed as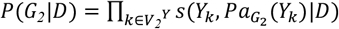, where *s* is the score function that only depends on a node (e.g. *Y*_*k*_) and its parent set (e.g. 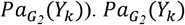 includes both ncRNA *X*’s and gene *Y*’s as the parent set of *Y*_*k*_ in 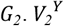 is the gene nodes in *V*_*2*_. Note that based on our semi-bipartite graph structure in Fig 1A, the ncRNA node *X*_*j*_ has the parent set 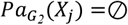 thus 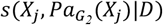 does not contribute to the final score and is not included.

The simplest approach is to do a global search for the DAG *G* with the highest BGe score. However, with a limited sample size, a single model usually provides poor approximation and the structure found is quite sensitive to the search procedure used. Alternatively, we can use MCMC simulation to sample a set of structures representative of the posterior to approximate Bayesian true model averaging. However, the space of DAG structures grows super-exponentially with the number of variables. Even with our reduced space after stage I, the computation is still not feasible. To remedy this, Friedman and Koller (2003) proposed a solution to sample over the space of orderings, called order MCMC^13^. We here proposed a new order MCMC algorithm adapted to our context by considering the semi-bipartite graph nature of NRN to further restrict the parent selection and speed up the computation, incorporating the prior knowledge during sampling to prioritize searching for validated features (e.g. nodes and edges) that are of more biological interest.

Denote by ≺ the topological order of nodes in a graph, and we associate a permutation *π*_≺_ with each order ≺. If a DAG *G* is compatible with an order ≺ or *G ∈≺*, the parents of each node must have a higher index in the permutation, i.e.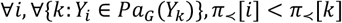. In a semi-bipartite graph structure, ncRNA nodes *X*’s are the parents of gene nodes *Y*’s, so *X*’s always have a higher index than their linked *Y*’s in the permutation. In order space, each order receives a score equal to the sum of the scores of all DAGs compatible with this order:

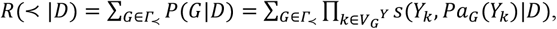

where *Γ*_≺_ denotes the set of all DAGs compatible with 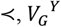 is the gene nodes in *G*. With the stage I screening, the order searching space has been greatly reduced. We will use MCMC to sample over the space of orderings and obtain the overall probability of a particular feature occurring. At each iteration t, we propose a move to change the order: (1) global swap, in which any two random nodes will swap positions in the node ordering while keeping other fixed; (2) local transposition, in which only two adjacent nodes will swap positions; or (3) node relocation, in which we place a single node in any position of the current order and accept with probability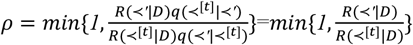, where ≺^[*t*]^ is the current ordering in t-th iteration and ≺ ′ the proposed new ordering. *q*(≺ ′| ≺^[*t*]^) is the proposal probability which defines the probability that the algorithm will “propose” a move from ≺^[*t*]^ to ≺ ′, and we usually assume equal proposal where *q*(≺ ′| ≺^[ *t*]^)=*q*(≺^[*t*]^ | ≺ ′) thus can be cancelled out. If accepted, then ≺^[ *t*+1]^ =≺ ′, otherwise ≺^[ *t*+1]^ = ≺^[ *t*]^. The resulting chain is reversible and has the desired stationary distribution^64^. We follow from Kuipers et al. (2022)^65^ to further perform more efficient sampling, the computational complexity of our algorithm has dropped to 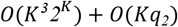, where K is the maximum size of potential parent sets.

The above order MCMC algorithm generates a chain of orders. We will then sample a DAG consistent with the sampled order by sampling the parents of each node proportionally to the entries respecting the order in the score table as in Kuipers et al. (2022)^65^. Assuming that the Markov chain has converged within T steps, and we obtain a sample of DAGs *G*^(1)^, …, *G*^(D)^, after excluding the first t’ burn-in steps, we can approximate the posterior probability of a specific structure feature *f* (e.g. one edge or a set of edges) by the sample average 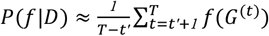, where 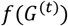 equals 1 if the feature is present in *G*^(*t*)^ and 0 otherwise. The final graph with the highest posterior probability can be identified accordingly^65^. Detailed algorithms can be found in the Supplementary section S1.5.

#### 1.3 Incorporation of prior knowledge from curated database

When inclusion and exclusion of all nodes and edges are equally alike, we will have the same probability for sampling each edge when drawing the DAG, i.e. the probability of sampling an edge from jth ncRNA to kth gene *P*(*e*_*jk*_) = *1*/*m* for *m* total possible edges that can be sampled. However, for ncRNAs and genes, scientists have conducted many animal or human studies to identify critical ncRNA-disease associations and ncRNA-gene interactions based on experiment, sequence (e.g. motif matching in miRNA-target gene searching) or additional biological information other than the transcriptomic data.

These information have been collected in the curated database and supplement the transcriptomic data (which could be restricted by sample size, condition, etc.) in revealing the underlying true regulatory network, ideally the regulatory network search algorithm should account for these contexts and prioritize the validated nodes or edges in the graph search. We have carefully preprocessed (download, annotate and categorize by diseases, etc.) and put together a pool of validated ncRNA-target gene edges as well as ncRNA nodes associated with diseases or conditions from existing curated databases on miRNAs, lncRNAs and other types of ncRNAs and their relationship with genes and diseases (see a full list in Table S1). When applying the algorithm to real data examples, we selected the subset of ncRNA nodes and ncRNA-gene interactions for the relevant disease or condition from the pool and incorporated as prior knowledge into our algorithm. We penalized the edges not belonging to the database by a factor of 2 thus those edges belonging to the database will have a relatively higher probability to be sampled. Suppose we have a collective database *Ω*_*G*_ for validated ncRNA->gene edges and a collective database *Ω*_*n*_ for disease associated ncRNAs. If the edge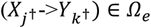, then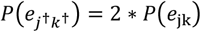, for *j* ≠ *j, k*^†^≠ *k*^†^ in MCMC sampling (all probabilities will be scaled so the sum of all probabilities is still equal to 1, so only the relative probabilities matter). If an ncRNA node 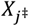 was found related to disease of interest in the database *Ω*_*n*_, i.e.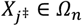, we penalize edges from the other ncRNA nodes so the edges with the ncRNA of interest as parent will have higher relative probability 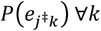to be sampled. This way, validated disease-associated ncRNA nodes and ncRNA-gene regulatory pairs were more likely to be sampled and remained in the final network.

### 2. Identifying differential ncRNA regulatory network

The above two-stage algorithm generates an optimal ncRNA-gene regulatory network based on the expression data of all samples under study. In real data, however, the expression data can be collected from multiple heterogeneous but comparable conditions (e.g. different clinical conditions, different tissues, different cell types, etc.). The ncRNA regulatory network could vary condition by condition implying the difference in underlying molecular regulatory mechanisms. For example in cancer study, researchers collect ncRNA and gene expression data from both early stage and late stage samples. The ncRNA regulatory network in late stage cancer can be different from that in early stage cancer implying the change in disease regulatory mechanism as the cancer progresses. In this paper, we borrowed the idea from incremental learning: we first used one condition as reference to generate an initial regulatory network by applying the two-stage hybrid method described above; then we updated the network using the data from the other condition by applying the proposed MCMC sampling scheme in stage II; this new network will be compared to the initial network for difference in connection patterns (e.g. what edges are gained or lost).

Suppose the samples are collected from two conditions (condition 1 vs 2, e.g. early stage vs late stage in cancer), a node ordering ≺_*cond 1*_ will first be learned from the expression data of ncRNA and gene from condition 1 (i.e. the reference condition) using the two-stage method. Next, we will run the order MCMC using data from condition 2 *D*_*cond2*_ to search for a global network of condition 2 from the initial network from condition 1. Suppose the network from condition 1 ends up with an ordering ≺_*cond 1*_, in the next iteration, we propose a new ordering ≺ ′ and update the MCMC chain with acceptance probability: *ρ* = 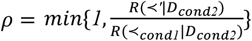. New moves (global swap, local transposition or node relocation) will be proposed in subsequent steps to generate a chain of orders and new DAGs for condition 2 can be sampled consistent with the order accordingly until convergence. This algorithm is fast and usually ends up with DAGs that are more comparable between two conditions. In the real data example, we applied the algorithm to compare DAG’s from early vs. late stage and identify how the ncRNA-gene regulatory relationship changes over different stages of cancer (differential regulation). These dynamic regulatory networks will potentially help provide insight into cancer progression. When the two conditions have imbalanced sample sizes, we found that the data with fewer samples (e.g. late stage cancer) tend to generate a more sparse network due to lack of power. To adjust for the difference that stems from imbalanced sample sizes in two conditions instead of the real biological difference in regulatory pattern, we reran the algorithm by updating the network using subsample from the condition with larger sample size and kept only those differential edges that are not differential in subsamples (Algorithm S6 in Supplementary). Note that the proposed algorithm is not invariant to the choice of the reference. We recommend using the one with large sample size as reference condition to construct a more robust and reliable initial network to start with and compare to.

### 3. Modularization of ncRNA regulatory network and pathway analysis

Most networks, including the ncRNA regulatory networks we study here, are found to divide naturally into communities or modules. The problem of detecting and characterizing this community structure can be in the study of networked systems. For better interpretation and more in-depth investigation of the network, we implement a directed Louvain algorithm^22^ to partition the final identified DAG into more interpretable modules and perform subnetwork analysis. The idea is that ncRNAs and genes within the same modules typically function together and provide more biological insight while nodes between modules are often loosely connected and their interaction is of less interest.

Depending on the biological questions of interest, we can perform downstream analysis on the identified network, e.g. network property analysis (density, clustering coefficient, identifying hub nodes or nodes with large betweenness), pathway analysis on the genes regulated by ncRNAs. For pathway analysis, we incorporate comprehensive pathway databases including KEGG^66^, Reactome^67^ and Biocarta^68^, the analysis can be performed both at a full network level or at module level.

### 4. R Shiny-based application

To run the complete pipeline of our framework and best visualize the resulting ncRNA regulatory network, we developed a R Shiny-based application with graphical user interface (GUI) (https://github.com/kehongjie/CAR-NET). This application took input of both ncRNA and gene expression data generated from either bulk or single-cell RNA-seq and incorporated a list of downloaded curated databases for users to choose based on the type of ncRNA and the condition/disease of interest. It starts with a preparatory step to preprocess the expression data by applying standard preprocessing for the different types of expression data. In its current version, *CAR-NET* only works on continuous data so standard normalization and transformation (log2 or other variance stabilizing transformations) should be applied for raw count data from bulk RNA-seq or scRNA-seq to generate continuous-valued data (e.g. log2 RPKM/FPKM/TPM) when necessary. For the best performance, we suggest filtering by means for bulk RNA-seq data, where the cutoff should be chosen looser for ncRNAs than coding genes due to their relatively low expression^69^. For scRNA-seq data, we also suggest filtering by percentage of zero counts across all cells for both ncRNAs and genes^70^. It then implements the major analytical steps to construct the ncRNA regulatory network, identify the differential network, detect network modules, and perform pathway analysis to facilitate the biological interpretation of the network findings. In addition, it provides visualization of the network/modules and downloadable graphical and tabular outputs. This GUI application runs a smoother and more standard pipeline with preset robust parameter values bypassing the technical and computationally intensive tuning processes, and provides an interactive interface for users to pick results of their interest to visualize (e.g. specific modules or specific ncRNA), is thus most suitable to researchers with little knowledge of programming and only basic statistical knowledge.

### 5. Comparative methods

To date, a number of methods have been developed to infer GRN from expression data and can be classified mainly in the following three categories^5-7^:

a. Information theory based methods (a.k.a. correlation-based methods): This category of methods typically use Pearson correlation or similar marginal measures to infer the regulatory network structure, are thus fast but can only generate undirected network. Representative methods include Pearson-correlation based method (marginal correlation hard threshold)^30^ and WGCNA (marginal correlation soft threshold)^31^. These methods typically apply hard threshold on correlation statistics or correlation test p-values after multiple comparison adjustment, or use soft-thresholding power to transform the correlation matrix into an adjacency matrix to construct the GRN. These methods are usually not computationally demanding thus appropriate to infer large GRNs, but the GRNs obtained by these methods are undirected networks.
b. Model based methods: This category of methods model gene expression profiles and then use the model to construct the network. It usually has high flexibility and scalability. Representative types of methods in this category include Bayesian network and differential equation based methods. These methods typically start by selecting a reasonable model to fit the data, followed by optimizing the model and constructing the GRN based on the optimized model. Model-based methods usually have high flexibility in modeling the different data types and network characteristics and incorporating prior biological knowledge but also have higher computational complexity.
c. Machine learning based methods: This category of methods uses machine learning calculation methods and data structures to fit gene expression data and reconstruct GRN. Representative methods in this category typically applied random forest (RF), boosting and other tree-based methods. These methods first transformed the problem into a classification or regression problem, then selected machine learning algorithm to target the problem, and build the network. Machine learning methods are suitable in modeling non-linear and dynamic behavior but also subject to high computational cost.

We chose 1-2 representative and computationally feasible methods from each of the above categories for comparison in case studies and simulations. For category (a), we chose Pearson-correlation based method (Pearson)^30^ and WGCNA^31^. For category (b), we compared our method to other popular Bayesian network structure learning methods including PC^28^, FCI^59^ and other hybrid algorithms. For category (c), we compared our method to a popular GRN tool that relies on random forest called GENIE3^29^.

### 6. Overview of simulation

Other than the three real data case studies, we also conducted extensive simulations with low-to-high dimension of ncRNA and gene expression data to compare our method to the popular GRN inferring methods. In brief, we found our method was more powered in removing irrelevant edges (thus a more sparse network) than other BN structure learning methods (e.g. PC, FCI) and GRN inferring methods (e.g. Pearson, GENIE3, WGCNA) while keeping a high sensitivity, i.e. most true edges were still kept (Extended Data Fig4 and Supplementary section S2.2), regardless of the cutoffs used in each method (we compared over a range of cutoffs for all methods). In addition, *CAR-NET* is computationally much faster than most other GRN inferring methods except for Pearson method (Table S11). Detailed simulation setting and main simulation results can be found in Supplementary section S2.

## Data availability

All datasets used in this work are publicly available. The lncRNA and gene expression data of human brain samples in case study 1 is available at LncExpDB database^26^. The miRNA and gene expression data of TCGA KIRP cohort in case study 2 is available at LinkedOmics^41^. The single-cell RNA-seq data from Smart-seq-total platform used in case study 3 is available in the GEO database under accession code GSE151334.

## Code availability

The code used to develop the model and generate results in this study as well as the software to implement the methods is publicly available and has been deposited in GitHub at: https://github.com/kehongjie/CAR-NET.

## Author Contributions

HK took the lead in developing the method, the software and wrote the manuscript. ZY, LF, ZX, RP, EL are involved in the development of the software. TM conceptualized the idea, supervised the project, took the lead in editing the manuscript, and acquired the funding. JQ, SC and ML contributed to manuscript writing and polishing. All authors provided critical feedback and helped to shape the research, analysis, and manuscript.

## Acknowledgements

Research reported in this publication was supported by the National Institute on Drug Abuse (NIDA) of National Institute of Health under the award number 1DP1DA048968 to SC and TM, by the National Institute on Drug Abuse (NIDA) under the award number 1K01DA059603 to TM, by the University of Maryland MPower Brain Health and Human Performance seed grant, Grand Challenge Grant, EPIB Department Pilot Award to TM.

## Additional Information

Extended Data Figures 1-4

## Supplementary Information

Supplementary methods and simulation: CAR-NET_supp.pdf

Supplementary tables: CAR-NET_supp_table.pdf

## Extended Data

**Extended Data Figure 1.**
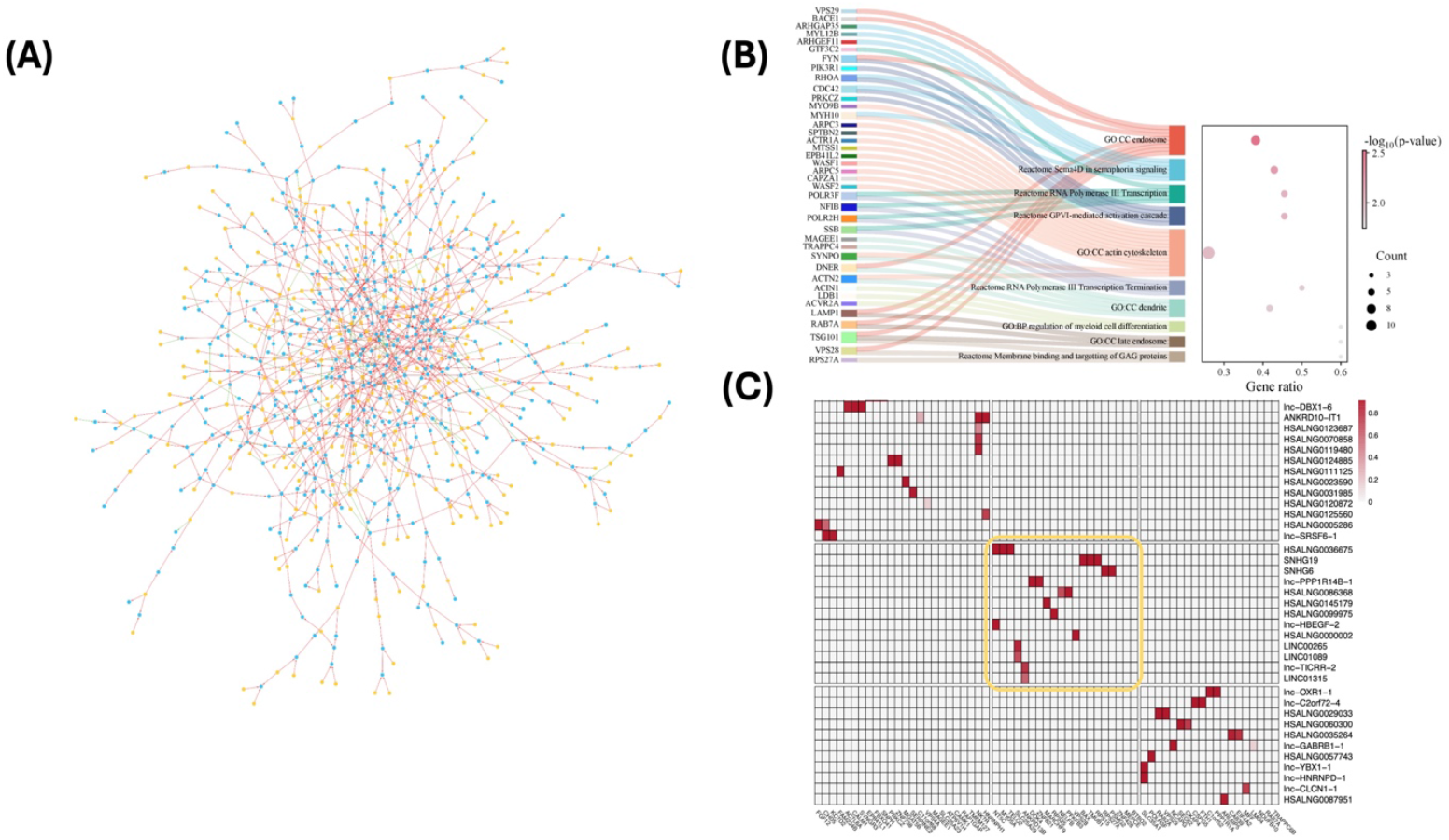
(A) The final lncRNA regulatory network detected by *CAR-NET* for case study 1: the brain development example. Yellow nodes refer to lncRNAs and blue nodes refer to genes. Red edges indicate activation and green edges indicate inhibition; (B) Sankey plot of the top 10 pathways enriched with genes from the final LncRNA regulatory network; (C) lncRNA by gene heatmap for the top three largest modules detected from the final ncRNA regulatory network. Each row indicates an lncRNA and each column indicates a gene. Color scale indicates the posterior probability of the corresponding edge from our method. Highlighted module corresponds to the one we show in Figure 3D of the main text.

**Extended Data Figure 2.**
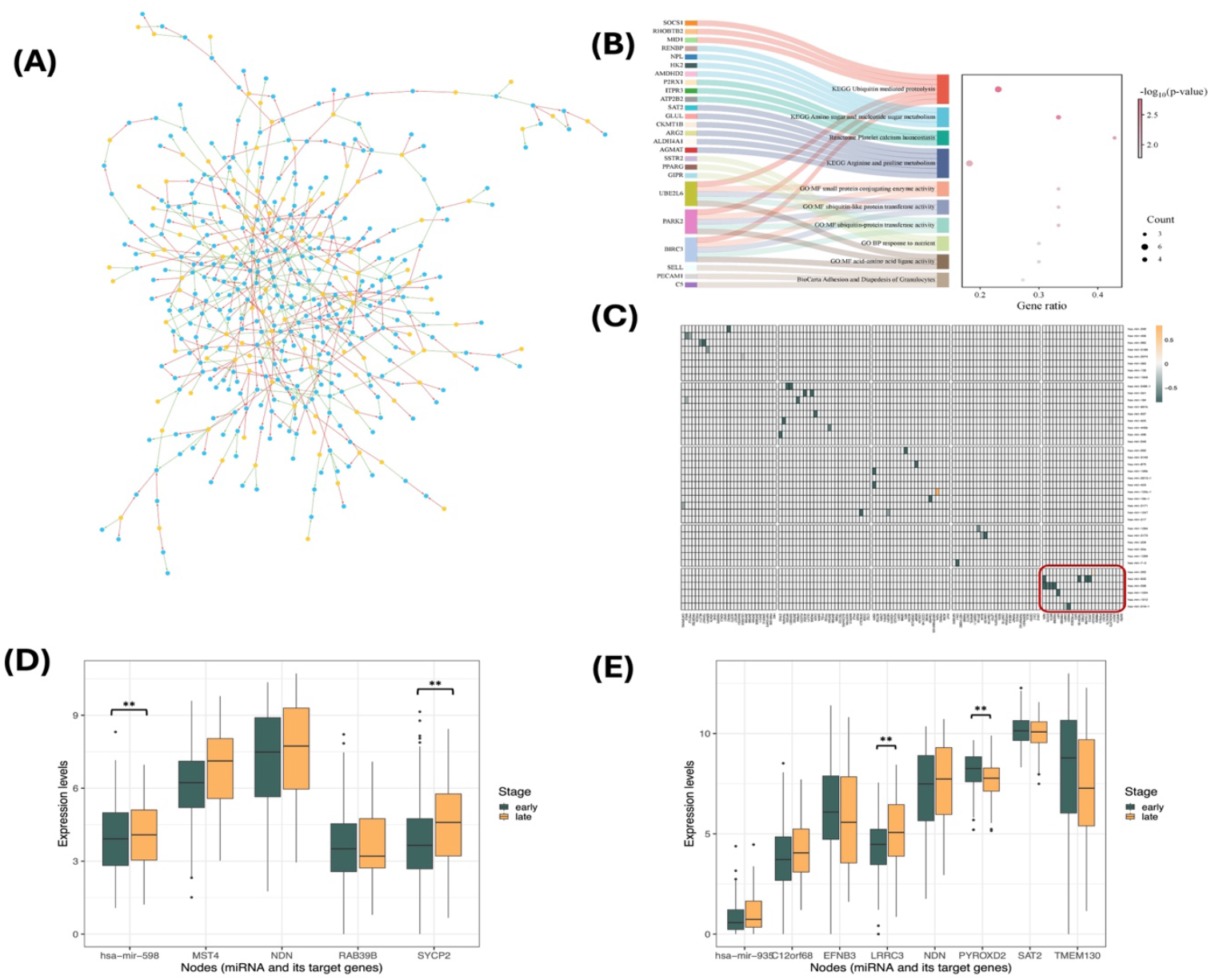
(A) The final miRNA regulatory network detected by *CAR-NET* from the early stage samples of KIRP in TCGA in case study 2. Yellow nodes refer to lncRNAs and blue nodes refer to genes. Red edges indicate activation and green edges indicate inhibition; (B) Sankey plot for the top 10 pathways enriched with genes in the final network generated from early stage KIRP samples; (C) lncRNA by gene heatmap for the top five largest modules differentially regulated between early and late stages after adjustment. Only differential edges that are not differential in subsamples are kept to reduce the bias introduced by imbalanced sample size. Each row indicates an miRNA and each column indicates a gene. Color scale indicates difference in the posterior probability (more yellowish/positive value means late stage has higher posterior probability while more greenish/negative value means late stage has lower posterior probability); (D) Box plots of expression levels for miRNA hsa-mir-598 and its regulated genes in KIRP early vs late stage. We use the following convention for symbols indicating statistical significance: * p<0.05; ** p<0.01; ***p<0.001. P-values are from two-sided t-tests; (E) Box plots of expression levels for miRNA hsa-mir-935 and its regulated genes in KIRP early vs late stage. We use the following convention for symbols indicating statistical significance: * p<0.05; ** p<0.01; ***p<0.001. P-values are from two-sided t-tests.

**Extended Data Figure 3.**
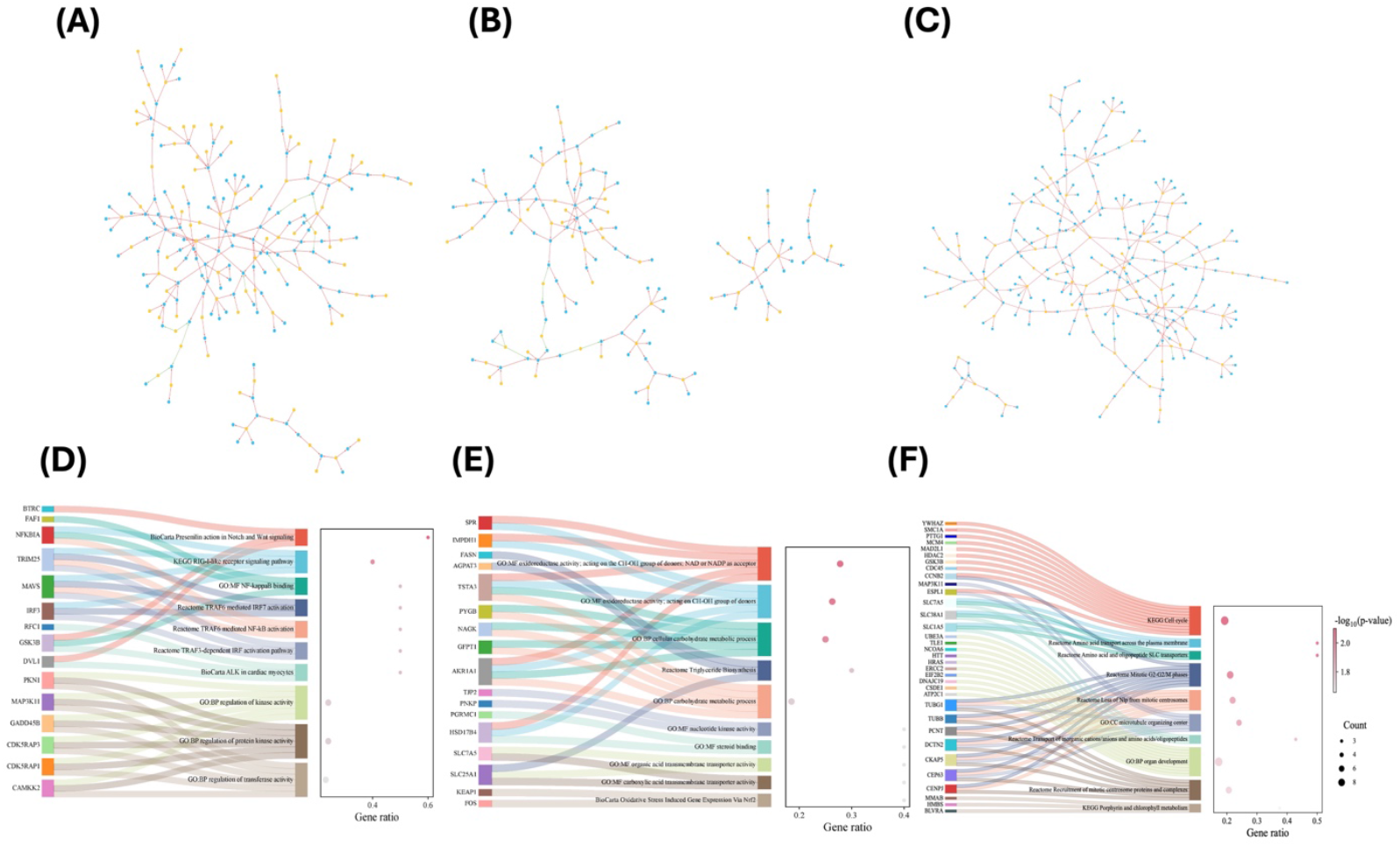
(A) Full ncRNA regulatory network for Fibroblast cell line in case study 3; (B) Full ncRNA regulatory network for HEK293 cell line in case study 3; (C) Full ncRNA regulatory network for MCF7 cell line in case study 3; (D) Sankey plot for the top 10 pathways enriched with genes in the final network for Fibroblast cell line; (E) Sankey plot for the top 10 pathways enriched with genes in the final network for HEK293 cell line; (F) Sankey plot for the top 10 pathways enriched with genes in the final network for MCF7 cell line.

**Extended Data Figure 4.**
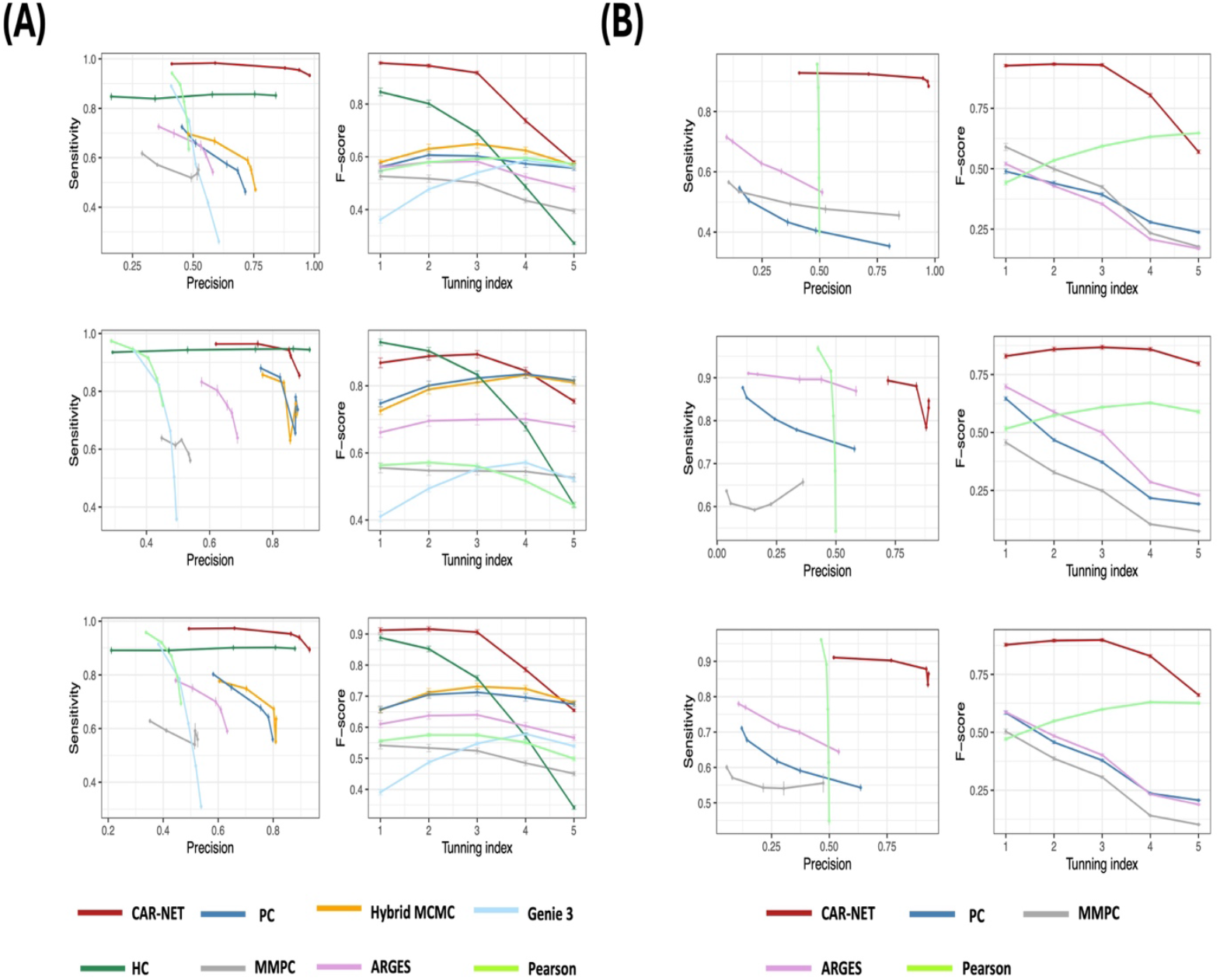
(A) Simulation results in low-dimension setting for level-1 edges (Upper), level-2 edges (Middle) and all edges (Lower). The left panel is the sensitivity-precision plot (a.k.a. precision-recall curve) and the right panel is comparison of F-score over varying thresholds/tuning indices for different methods; (A) Simulation results in high-dimension setting for level-1 edges (Upper), level-2 edges (Middle) and all edges (Lower). The left panel is the sensitivity-precision plot (a.k.a. precision-recall curve) and the right panel is comparison of F-score over varying thresholds/tuning indices for different methods. Methods running into computational bottleneck in high-dimensional case are not included for comparison.

## References

1 Kaikkonen, M. U., Lam, M. T. & Glass, C. K. Non-coding RNAs as regulators of gene expression and epigenetics. Cardiovascular research 90, 430–440 (2011).

2 Statello, L., Guo, C.-J., Chen, L.-L. & Huarte, M. Gene regulation by long non-coding RNAs and its biological functions. Nature reviews Molecular cell biology 22, 96–118 (2021).

3 Esteller, M. Non-coding RNAs in human disease. Nature Reviews Genetics 12, 861–874 (2011). 10.1038/nrg3074

4 Chai, L. E. et al. A review on the computational approaches for gene regulatory network construction. Computers in biology and medicine 48, 55–65 (2014).

5 Badia-i-Mompel, P. et al. Gene regulatory network inference in the era of single-cell multi-omics. Nature Reviews Genetics 24, 739–754 (2023).

6 Zhao, M., He, W., Tang, J., Zou, Q. & Guo, F. A comprehensive overview and critical evaluation of gene regulatory network inference technologies. Briefings in bioinformatics 22, bbab009 (2021).

7 Delgado, F. M. & Gómez-Vela, F. Computational methods for gene regulatory networks reconstruction and analysis: a review. Artificial intelligence in medicine 95, 133–145 (2019).

8 Anastasiadou, E., Jacob, L. S. & Slack, F. J. Non-coding RNA networks in cancer. Nature Reviews Cancer 18, 5–18 (2018).

9 Van Steen, K. Travelling the world of gene–gene interactions. Briefings in bioinformatics 13, 1–19 (2012).

10 Panni, S., Lovering, R. C., Porras, P. & Orchard, S. Non-coding RNA regulatory networks. Biochimica et biophysica acta (BBA)-Gene regulatory mechanisms 1863, 194417 (2020).

11 Xu, J. et al. The mRNA related ceRNA–ceRNA landscape and significance across 20 major cancer types. Nucleic acids research 43, 8169–8182 (2015).

12 Leclerc, R. D. Survival of the sparsest: robust gene networks are parsimonious. Molecular systems biology 4, 213 (2008).

13 Friedman, N. & Koller, D. Being Bayesian about network structure. A Bayesian approach to structure discovery in Bayesian networks. Machine learning 50, 95–125 (2003).

14 Huang, H.-Y. et al. miRTarBase 2020: updates to the experimentally validated microRNA–target interaction database. Nucleic acids research 48, D148–D154 (2020).

15 L, C. et al. LncRNA2Target v2.0: a comprehensive database for target genes of lncRNAs in human and mouse. Nucleic acids research 47 (2019). 10.1093/nar/gky1051

16 Zhou, B. et al. EVLncRNAs 3.0: an updated comprehensive database for manually curated functional long non-coding RNAs validated by low-throughput experiments. Nucleic Acids Research 52, D98–D106 (2024).

17 B, X., Q, D., H, H. & D, W. miRCancer: a microRNA-cancer association database constructed by text mining on literature. Bioinformatics (Oxford, England) 29 (2013). 10.1093/bioinformatics/btt014

18 Bao, Z. et al. LncRNADisease 2.0: an updated database of long non-coding RNA-associated diseases. Nucleic acids research 47, D1034–D1037 (2019).

19 Michoel, T. & Nachtergaele, B. Alignment and integration of complex networks by hypergraph-based spectral clustering. Physical Review E—Statistical, Nonlinear, and Soft Matter Physics 86, 056111 (2012).

20 Grimes, T., Potter, S. S. & Datta, S. Integrating gene regulatory pathways into differential network analysis of gene expression data. Scientific reports 9, 5479 (2019).

21 Van de Ven, G. M., Tuytelaars, T. & Tolias, A. S. Three types of incremental learning. Nature Machine Intelligence 4, 1185–1197 (2022).

22 Blondel, V. D., Guillaume, J.-L., Lambiotte, R. & Lefebvre, E. Fast unfolding of communities in large networks. Journal of statistical mechanics: theory and experiment 2008, P10008 (2008).

23 H, Z. et al. LncTarD 2.0: an updated comprehensive database for experimentally-supported functional lncRNA-target regulations in human diseases. Nucleic acids research 51 (2023). 10.1093/nar/gkac984

24 Wu, P. et al. Roles of long noncoding RNAs in brain development, functional diversification and neurodegenerative diseases. Brain research bulletin 97, 69–80 (2013).

25 Aliperti, V., Skonieczna, J. & Cerase, A. Long non-coding RNA (lncRNA) roles in cell biology, neurodevelopment and neurological disorders. Non-coding RNA 7, 36 (2021).

26 Li, Z. et al. LncExpDB: an expression database of human long non-coding RNAs. Nucleic Acids Research 49, D962–D968 (2021).

27 Sarropoulos, I., Marin, R., Cardoso-Moreira, M. & Kaessmann, H. Developmental dynamics of lncRNAs across mammalian organs and species. Nature 571, 510–514 (2019).

28 Kalisch, M. & Bühlman, P. Estimating high-dimensional directed acyclic graphs with the PC-algorithm. Journal of Machine Learning Research 8 (2007).

29 Huynh-Thu, V. A., Irrthum, A., Wehenkel, L. & Geurts, P. Inferring regulatory networks from expression data using tree-based methods. PloS one 5, e12776 (2010).

30 Song, L., Langfelder, P. & Horvath, S. Comparison of co-expression measures: mutual information, correlation, and model based indices. BMC bioinformatics 13, 1–21 (2012).

31 Langfelder, P. & Horvath, S. WGCNA: an R package for weighted correlation network analysis. BMC bioinformatics 9, 1–13 (2008).

32 Yap, C. C. & Winckler, B. Harnessing the power of the endosome to regulate neural development. Neuron 74, 440–451 (2012).

33 Jan, Y.-N. & Jan, L. Y. The control of dendrite development. Neuron 40, 229–242 (2003).

34 Ransohoff, R. M. & Cardona, A. E. The myeloid cells of the central nervous system parenchyma. Nature 468, 253–262 (2010).

35 Hetman, M. & Slomnicki, L. P. Ribosomal biogenesis as an emerging target of neurodevelopmental pathologies. Journal of neurochemistry 148, 325–347 (2019).

36 Young, C. et al. Ethanol-induced neuronal apoptosis in vivo requires BAX in the developing mouse brain. Cell Death & Differentiation 10, 1148–1155 (2003).

37 Andersen, S. L. Trajectories of brain development: point of vulnerability or window of opportunity? Neuroscience & Biobehavioral Reviews 27, 3–18 (2003).

38 Gil, N. & Ulitsky, I. Regulation of gene expression by cis-acting long non-coding RNAs. Nature Reviews Genetics 21, 102–117 (2020).

39 Network, C. G. A. R. Comprehensive molecular characterization of papillary renal-cell carcinoma. New England Journal of Medicine 374, 135–145 (2016).

40 Munari, E. et al. Clear cell papillary renal cell carcinoma: micro-RNA expression profiling and comparison with clear cell renal cell carcinoma and papillary renal cell carcinoma. Human pathology 45, 1130–1138 (2014).

41 SV, V., P, S., J, W. & B, Z. LinkedOmics: analyzing multi-omics data within and across 32 cancer types. Nucleic acids research 46 (2018). 10.1093/nar/gkx1090

42 Lin, S. & Gregory, R. I. MicroRNA biogenesis pathways in cancer. Nature reviews cancer 15, 321–333 (2015).

43 Peng, Y. & Croce, C. M. The role of MicroRNAs in human cancer. Signal transduction and targeted therapy 1, 1–9 (2016).

44 De Faveri, L. et al. Putative tumour suppressor gene necdin is hypermethylated and mutated in human cancer. British journal of cancer 108, 1368–1377 (2013).

45 Schmit, K. & Michiels, C. TMEM proteins in cancer: a review. Frontiers in pharmacology 9, 1345 (2018).

46 Jovic, D. et al. Single-cell RNA sequencing technologies and applications: A brief overview. Clinical and translational medicine 12, e694 (2022).

47 Isakova, A., Neff, N. & Quake, S. R. Single-cell quantification of a broad RNA spectrum reveals unique noncoding patterns associated with cell types and states. Proceedings of the National Academy of Sciences 118, e2113568118 (2021).

48 Zhang, X., Hamblin, M. H. & Yin, K.-J. The long noncoding RNA Malat1: Its physiological and pathophysiological functions. RNA biology 14, 1705–1714 (2017).

49 Canida, T., Ke, H., Chen, S., Ye, Z. & Ma, T. Multivariate Bayesian variable selection for multi-trait genetic fine mapping. Journal of the Royal Statistical Society Series C: Applied Statistics, qlae055 (2024).

50 Jumper, J. et al. Highly accurate protein structure prediction with AlphaFold. nature 596, 583–589 (2021).

51 Pearl, J. Causal inference in statistics: An overview. (2009).

52 Koller, D. (The MIT Press, 2009).

53 J, F. & J, L. Sure Independence Screening for Ultrahigh Dimensional Feature Space. Journal of the Royal Statistical Society Series B: Statistical Methodology 70, 849–911 (2008). 10.1111/j.1467-9868.2008.00674.x

54 Bühlmann, P., Kalisch, M. & Maathuis, M. H. Variable selection in high-dimensional linear models: partially faithful distributions and the PC-simple algorithm. Biometrika 97, 261–278 (2010).

55 Ke, H. et al. High-dimension to high-dimension screening for detecting genome-wide epigenetic and noncoding RNA regulators of gene expression. Bioinformatics 38, 4078–4087 (2022). 10.1093/bioinformatics/btac518

56 Ma, T., Ke, H. & Ren, Z. Robust distance correlation for variable screening. arXiv preprint arXiv:2212.13292 (2022).

57 Zhu, A., Ibrahim, J. G. & Love, M. I. Heavy-tailed prior distributions for sequence count data: removing the noise and preserving large differences. Bioinformatics 35, 2084–2092 (2019).

58 Ke, Y., Minsker, S., Ren, Z., Sun, Q. & Zhou, W.-X. User-friendly covariance estimation for heavy-tailed distributions. Statistical Science 34, 454–471 (2019).

59 Spirtes, P. L., Meek, C. & Richardson, T. S. Causal inference in the presence of latent variables and selection bias. arXiv preprint arXiv:1302.4983 (2013).

60 Cohen, J. Statistical power analysis for the behavioral sciences. (routledge, 2013).

61 Velickovic, P. et al. Graph attention networks. stat 1050, 10–48550 (2017).

62 Wille, A. & Bühlmann, P. Low-order conditional independence graphs for inferring genetic networks. Statistical applications in genetics and molecular biology 5 (2006).

63 Wilks, S. The likelihood test of independence in contingency tables. The Annals of Mathematical Statistics 6, 190–196 (1935).

64 Gilks, W. R., Richardson, S. & Spiegelhalter, D. Markov chain Monte Carlo in practice. (CRC press, 1995).

65 Kuipers, J., Suter, P. & Moffa, G. Efficient sampling and structure learning of Bayesian networks. Journal of Computational and Graphical Statistics 31, 639–650 (2022).

66 Kanehisa, M., Sato, Y., Kawashima, M., Furumichi, M. & Tanabe, M. KEGG as a reference resource for gene and protein annotation. Nucleic acids research 44, D457–D462 (2016).

67 Fabregat, A. et al. The reactome pathway knowledgebase. Nucleic acids research 46, D649–D655 (2018).

68 Nishimura, D. BioCarta. Biotech Software & Internet Report: The Computer Software Journal for Scient 2, 117–120 (2001).

69 Mattick, J. S. et al. Long non-coding RNAs: definitions, functions, challenges and recommendations. Nature reviews Molecular cell biology 24, 430–447 (2023).

70 Heumos, L. et al. Best practices for single-cell analysis across modalities. Nature Reviews Genetics 24, 550–572 (2023).

