## Supplementary methods for "Inferring non-coding RNA regulatory network from transcriptomic data and curated databases"

March 9, 2025

### S1 Supplementary methods

#### S1.1 Overview of Bayesian network structure learning methods

Bayesian network (BN) structure learning is computationally challenging, to date a majority of methods that have been developed fall into three main categories (Pearl, 2009; Koller and Friedman, 2009): 1) constraint-based approaches: this category of methods, including the famous PC (Kalisch and Bühlman, 2007) and FCI algorithms (Spirtes et al., 2013), views BN as a representation of dependencies and tries to perform a series of conditional independent tests to find a network that best explains these dependencies. However, these methods are sensitive to failures in individual independent tests and error propagation thus are less accurate; 2) score-based approaches: this category of methods views BN as specifying a statistical model and addresses learning as a model selection problem. Representative methods under this category include HC (Heckerman et al., 1995), GIES (Hauser and Bühlmann, 2012) and GES (Chickering, 2002). They typically define a score function (mostly likelihood-based, e.g. BIC) that measures how well the model fits the observed data and then find the BN that maximizes the score. However, the number of BN structures to be searched is super-exponential in the number of nodes, in our context typically both ncRNA and gene expression data are high-dimensional, i.e. $p \gg n$ and $q \gg n$, making this category of methods computationally infeasible to directly detect the regulatory network; 3) hybrid approaches: this category of methods, usually implemented in two stages, combine the strengths of the previous two categories by implementing constraint-based strategy in the first stage to reduce the space of candidate DAGs, and a score-based strategy in the second stage to find the optimal DAG. Representative hybrid approaches include MMHC (Tsamardinos et al., 2006) and ARGES (Nandy et al., 2018). However, none of the existing hybrid approaches are specific to construct the ncRNA-gene regulatory network with unique structure in our study.

#### S1.2 BN structure learning stage I: edge-wise screening

Similar to existing constraint-based approaches (Kalisch and Bühlman, 2007), our method also relies on conditional independence test to perform edge-wise screening to remove implausible edges in stage I. We have previously proposed the robust partial correlation statistics (rPCor) for screening in multivariate linear regression to obtain a sparse regression coefficient matrix with theoretical guarantee (Ke et al., 2022; Ma et al., 2022). As both multivariate linear regression and Gaussian Bayesian network assumed conditional independence and structural sparsity (Chiquet et al., 2017; Vogels et al., 2024), it becomes natural to adopt the rPCor statistics and extend its use for edge-wise screening in BN (see the connection in section S1.7). The rPCor statistics we used to perform conditional independence tests to remove irrelevant level 1 and level 2 edges are $\rho_\tau(X_j, Y_k|S_{X_j}, S_{Y_k})$ and $\rho_\tau(Y_l, Y_{l'}|S_{Y_l}, S_{Y_{l'}})$, where $S_{X_j}$, $S_{Y_k}$, $S_{Y_l}$ and $S_{Y_{l'}}$ are the conditional sets of nodes ($S_{X_j}$ include ncRNA nodes and the other three sets include gene nodes). In this study, we will adopt a modified version of rPCor statistics with carefully selected conditional sets, determine the optimal test threshold in the network structure learning context, and iteratively perform conditional independence tests on rPCor to remove implausible ncRNA-gene and gene-gene edges.

##### S1.2.1 Selection of conditional set

Partial correlation-based methods typically search over all nodes in the full conditional set when testing for the partial correlation, which is computationally heavy. In ncRNA-gene regulation problem, we know a majority of ncRNA nodes are unrelated and most genes are unrelated over the whole genome. We propose to apply a marginal correlation threshold and regard only those ncRNAs/genes having marginal correlation above certain threshold (e.g. $|\rho| > 0.3$ as in Cohen's guideline (Cohen, 2013)) with the nodes of interest as the conditional sets. For example for level-1 ncRNA-gene edges, we can find the neighboring set $\mathcal{M}(X_j) = \{j' : |\hat{\rho}(X_j, X_{j'})| > 0.3\} \setminus \{j\}$ for each $X_j$ and the neighboring set $\mathcal{M}(Y_k) = \{k' : |\hat{\rho}(Y_k, Y_{k'})| > 0.3\} \setminus \{k\}$ for each $Y_k$. The same idea has been used in masked attention in Graphical Attention Network (Veličković et al., 2017). Additionally, one can also consider using the Markov blanket to select the conditional set (Pearl, 2014). Unlike the conditional independence test threshold $\alpha$ directly used to select edges, the edge screening results are not too sensitive to the choice of the marginal correlation threshold used here for conditional set selection. The main purpose is to speed up the computation when screening for edges. To further improve the computation, as in (Bühlmann et al., 2010; Ke et al., 2022) we iteratively increase the size of the conditional set while performing edge screening.

##### S1.2.2 Selection of optimal conditional independence test threshold $\alpha$

The choice of conditional independence test threshold $\alpha$ is critical as it balances between the sensitivity and false positive control of the method. If $\alpha$ is chosen too small (too stringent cutoff), it is possible that important edges will be accidentally screened out. On the other hand, if $\alpha$ is chosen too large (too loose cutoff), most edges are kept so the network remains too dense. A majority of constraint-based approaches applied the conventional choice of 0.05 without much justification. We proposed a fast data-driven procedure using a pseudo F-score to select the optimal choice of $\alpha$. The regular F-score is defined as $F = \frac{2*precision*sensitivity}{precision+sensitivity}$. As we do not know the ground truth in real data, a pseudo F-score is defined with the same formula but uses a pseudo sensitivity and a pseudo precision, where the true edges are defined as those pairs with largest marginal correlations, to balance between effective dimension reduction (i.e. maintaining high precision) and high sensitivity in the remaining edges. Details procedure can be found in Algorithm S1. Our procedure can serve as an alternative to the slower stability selection-based procedure we have proposed before for screening in multivariate regression (Ke et al., 2022). In practice, we recommend users try out both to get the best results.

All the above have been put together to form our edge-wise screening algorithms to remove implausible level-1 (Algorithm S2) and level-2 edges (Algorithm S3).

**Algorithm S1:** A fast data-driven procedure to determine the optimal $\alpha$ value in stage I edge-wise screening.

Choose a sequence of candidate $\alpha$ values $(\alpha_1, \alpha_2, ..., \alpha_K)$ (e.g. (1e-3, 1e-4, 1e-5, 1e-6) with equal step size or with varying step). Here $\alpha_1$ is the largest value and $\alpha_K$ is the smallest value.

**Step 1.** For both ncRNA and gene expression data, we only keep the $p' = \lceil r * p \rceil$ ncRNAs and $q' = \lceil r * q \rceil$ genes with the largest variance and generate a sub-data. These ncRNAs and genes are the most informative ones so it's reasonable to infer the optimal $\alpha$ only based on this sub-data with $p'$ ncRNAs and $q'$ genes. We performed sensitivity analysis and the final $\alpha$ selected is not too sensitive to the choice of $r$, so we used $r = 0.2$ in all cases for the best computational efficiency.

**Step 2.** Calculate the marginal correlation between each ncRNA and gene, i.e. $\rho(X_j, Y_k)$ in the sub-data. Regard the top $\eta = \min\{p', q'\}$ ncRNA-gene pairs with the largest marginal correlation as "true" edges.

**Step 3.** Run the edge-wise screening using each value of $(\alpha_1, \alpha_2, ..., \alpha_K)$. Calculate the pseudo-sensitivity (=Number of "true" edges left in the network /Total number of "true" edges) and pseudo-precision (=Number of "true" edges left in the network/Total number of ncRNA-gene edges in the network), and pseudo-F= $\frac{2*pseudo-precision*pseudo-sensitivity}{pseudo-precision+pseudo-sensitivity}$ for each level of $\alpha$.

**Step 4.** Instead of using the absolute pseudo F-score to select the best $\alpha$, we learnt from the idea of elbow plot and gap statistics and considered to use the relative change in pseudo F-score to select the best $\alpha$, i.e. For $(F_{\alpha_1}, F_{\alpha_2}, ..., F_{\alpha_K})$ and chose the largest $\alpha$ where pseudo F-score starts to have a steep decline, e.g. the elbow point.

Figure S1 shows a demonstration of the optimal threshold $\alpha$ selection in KIRP miRNA-gene regulation example.

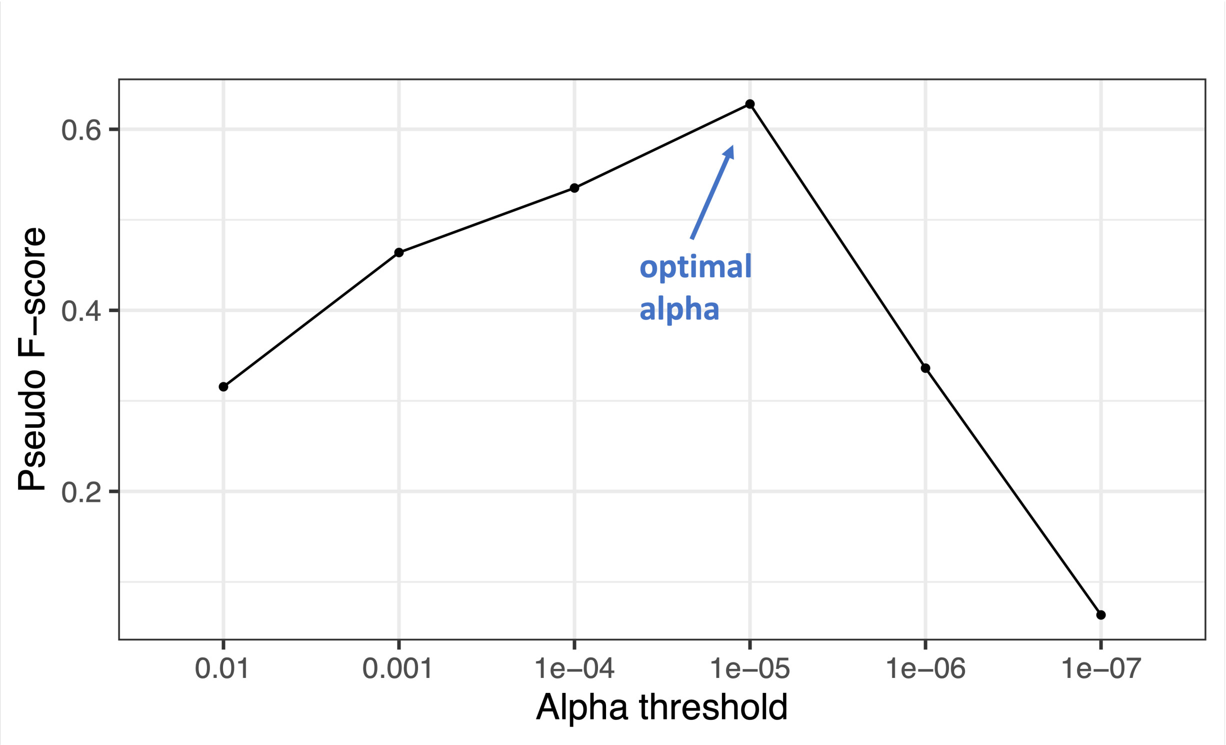


Figure S1: Tuning parameter $\alpha$ selection in KIRP miRNA network construction.

---

**Algorithm S2:** Edge-wise screening for level-1 edges

---

**Input:** ncRNA expression data $\boldsymbol{X} = (X_1, ..., X_p)$ and gene expression data $\boldsymbol{Y} = (Y_1, ..., Y_q)$, pre-determined optimal $\alpha$ from Algorithm S1

For each $X_j$, find the neighboring set $\mathcal{M}(X_j) = \{j' : |\hat{\rho}(X_j, X_{j'})| > 0.3\} \setminus \{j\}$;

For each $Y_k$, find the neighboring set $\mathcal{M}(Y_k) = \{k' : |\hat{\rho}(Y_k, Y_{k'})| > 0.3\} \setminus \{k\}$;

Suppose $m$ corresponds to the order of the conditional independence test, we start with $m = 1$ i.e. marginal screening, and build the active set for level-1 edges $\hat{\mathcal{A}_{xy}}^{[1]} = \{(j,k) : |(n-3)^{1/2} Z_\tau(X_j, Y_k)| > \Phi^{-1}(1-\alpha/2)\}$ where $Z_\tau(X_j, Y_k) = \frac{1}{2}\log(\frac{1+\rho_\tau(X_j,Y_k)}{1-\rho_\tau(X_j,Y_k)})$ is the Fisher z-transformation of correlation;

**repeat**

  $m = m + 1$;

  $\hat{\mathcal{A}_{xy}}^{[m]} = \hat{\mathcal{A}_{xy}}^{[m-1]}$;

  **repeat**

    Select a (new) pair of $(j,k) \in \hat{\mathcal{A}_{xy}}^{[m-1]}$;

    **if** $(n - |S_{X_j}| - |S_{Y_k}| - 3)^{1/2} |Z_\tau(X_j, Y_k \mid S_{X_j}, S_{Y_k})| \leq \Phi^{-1}(1-\alpha/2)$ *for any* $S_{X_j} \subseteq \Big\{\{j : (j,k) \in \hat{\mathcal{A}_{xy}}^{[m-1]}\} \cap \mathcal{M}(X_j)\Big\}$ *and* $S_{Y_k} \subseteq \Big\{\{k : (j,k) \in \hat{\mathcal{A}_{xy}}^{[m-1]}\} \cap \mathcal{M}(Y_k)\Big\}$ *with* $|S_{X_j}| + |S_{Y_k}| = m - 1$ **then**

      remove pair $(j,k)$ from $\hat{\mathcal{A}_{xy}}^{[m]}$

  **until** *all* $(j,k) \in \hat{\mathcal{A}_{xy}}^{[m-1]}$ *is tested*;

**until** $|\hat{\mathcal{A}_{xy}}^{[m]}| \leq m$ *or* $m = m_{max} = 5$;

**Output:** $\hat{\mathcal{A}_{xy}}^{[m]}$: the active set for remaining level-1 edges.

---

**Algorithm S3:** Edge-wise screening for level-2 edges

**Input:** Gene expression data $\boldsymbol{Y} = (Y_1, ..., Y_q)$, pre-determined $\alpha$ from algorithm S1

For each $Y_l$, find the neighboring set $\mathcal{M}(Y_l) = \{l' : |\hat{\rho}(Y_l, Y_{l'})| > 0.3\} \setminus \{l\}$;

As for level-1 edges, we start with $m = 1$ marginal screening and build the active set for level-2 edges $\hat{\mathcal{A}_{yy}}^{[1]} = \{(l, l') : |(n-3)^{1/2} Z_\tau(Y_l, Y_{l'})| > \Phi^{-1}(1 - \alpha/2)\}$ where $Z_\tau(Y_l, Y_{l'}) = \frac{1}{2}\log(\frac{1+\rho_\tau(Y_l,Y_{l'})}{1-\rho_\tau(Y_l,Y_{l'})})$ is the Fisher z-transformation of correlation;

**repeat**

$m = m + 1$;

$\hat{\mathcal{A}_{yy}}^{[m]} = \hat{\mathcal{A}_{yy}}^{[m-1]}$;

**repeat**

Select a (new) pair of $(l, l') \in \hat{\mathcal{A}_{yy}}^{[m-1]}$;

**if** $(n - |S_{Y_l}| - |S_{Y_l'}| - 3)^{1/2} |Z_\tau(Y_l, Y_{l'} \mid S_{Y_l}, S_{Y_l'})| \leq \Phi^{-1}(1 - \alpha/2)$ *for any* $S_{Y_l} \subseteq \left\{\{l : (l, l') \in \hat{\mathcal{A}_{yy}}^{[m-1]}\} \cap \mathcal{M}(Y_l)\right\}$ *and* $S_{Y_l'} \subseteq \left\{\{l' : (l, l') \in \hat{\mathcal{A}_{xy}}^{[m-1]}\} \cap \mathcal{M}(Y_l')\right\}$ *with* $|S_{Y_l}| + |S_{Y_l'}| = m - 1$ **then**

remove pair $(l, l')$ from $\hat{\mathcal{A}_{yy}}^{[m]}$

**until** *all* $(l, l') \in \hat{\mathcal{A}_{yy}}^{[m-1]}$ *is tested*;

**until** $|\hat{\mathcal{A}_{yy}}^{[m]}| \leq m$ *or* $m = m_{max} = 5$;

**Output:** $\hat{\mathcal{A}_{yy}}^{[m]}$: the active set for remaining level-2 edges.

#### S1.3 Orientation rules for level-2 edges after edge-wise screening

After edge-wise screening, we will identify the colliders and follow a few orientation rules to determine the directions of level-2 edges as in PC algorithm (Spirtes et al., 2001), e.g. orientation of v-structures (collider triples), avoiding new colliders, orientation to prevent cycles, etc.

In addition to existing orientation rules, the unique semi-bipartite graph structure of noncoding RNA regulatory network (NRN) enabled us to further orient the directions of level-2 edges. Suppose A is a ncRNA regulator, B is its immediate target gene and gene C is connected to gene B. A is independent of C conditional on B so no direct link exists between A and C. Suppose we do not know the direction of A-B edge, it will result in DAGs in Markov equivalent class (Figure S2 upper). With the direction of edge A $\rightarrow$ B given in the semi-bipartite graph (Fig 1A), we can break the Markov equivalence and orient B$\rightarrow$C (Fig S2 lower).

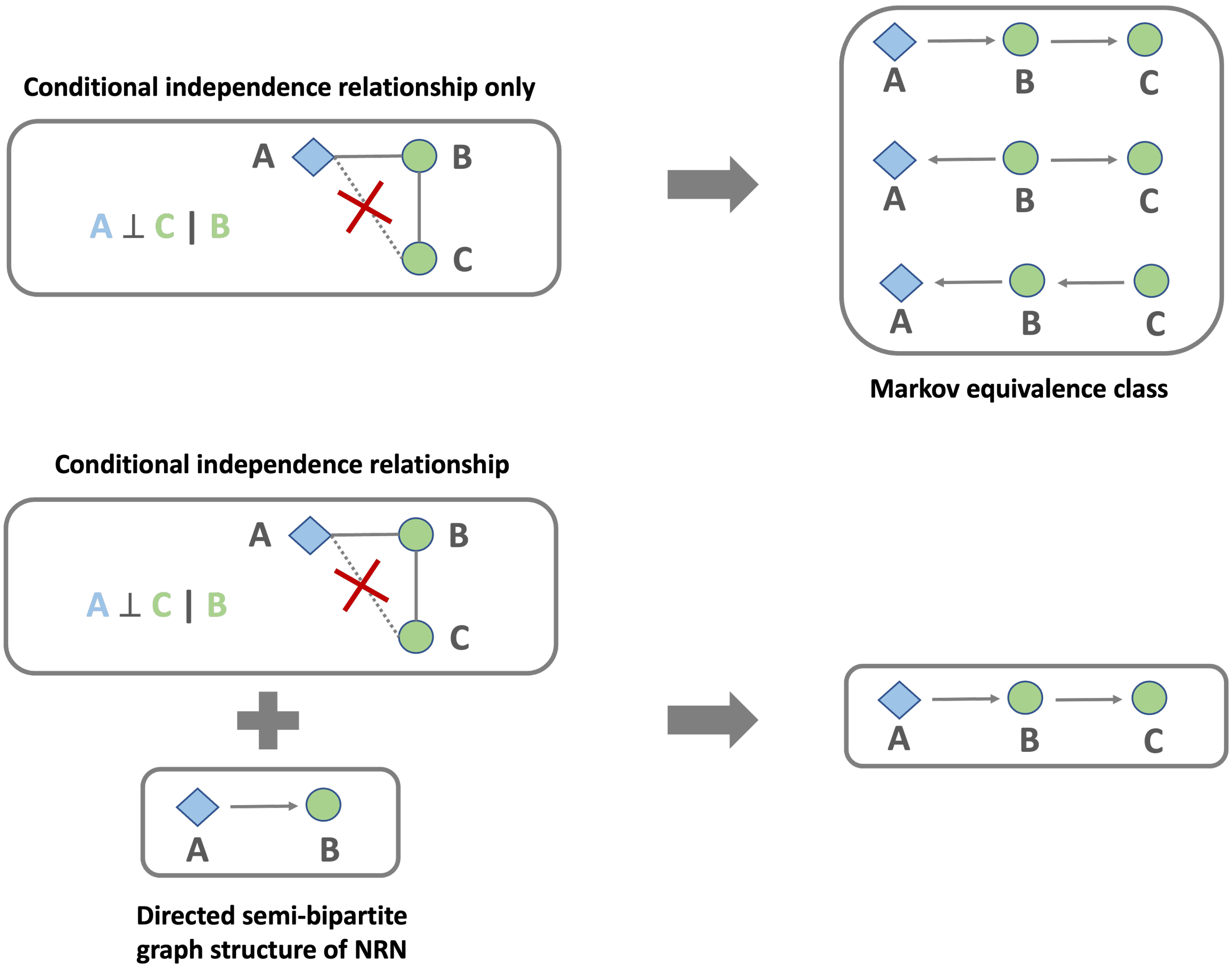


Figure S2: A toy example to show how the unique semi-bipartite graph structure of NRN helps us further orient the directions of level-2 edges.

#### S1.4 BN structure learning stage I: node-wise screening

After edge-wise screening (assume a partially directed graph $\mathcal{G}_1 = \{\mathcal{V}_1, \mathcal{E}_1\}$ left after edge-wise screening), it is common to see some ncRNA or gene nodes had very few edges connected (i.e. very small degree) left. These ncRNAs might be isolated (regulated genes not linked to the main network) or at the terminal of a network and thus were less critical to the full regulatory network. To guarantee more efficient computation and better convergence in the next stage, we further considered a node-wise screening by calculating a canonical correlation between each node and its remaining edges. For example, for every ncRNA $X_j$ left after the edge-wise screening, we compute its canonical correlation versus all the genes connected to this ncRNA (denoted as $\mathcal{N}(X_j)$) as:

$$\phi_j^X = \max_a \frac{\Sigma_{X_j \mathcal{N}(X_j)} a}{\sqrt{\Sigma_{X_j}} \sqrt{a^T \Sigma_{\mathcal{N}(X_j)} a}},$$

where $\mathcal{N}(X_j)$ includes all the gene nodes linked to jth ncRNA. $\Sigma_{X_j \mathcal{N}(X_j)}$ is the vector of covariance between ncRNA $X_j$ and its directly linked gene nodes. $\Sigma_{X_j}$ and $\Sigma_{\mathcal{N}(X_j)}$ are the variance of $X_j$ and the diagonal matrix of variances of its directly linked nodes.

Similarly, for each gene $Y_k$ left after the edge-wise screening, we can compute

$$\phi_k^Y = \max_b \frac{\Sigma_{Y_k \mathcal{N}(Y_k)} b}{\sqrt{\Sigma_{Y_k}} \sqrt{b^T \Sigma_{\mathcal{N}(Y_k)} b}},$$

where $\mathcal{N}(Y_k)$ includes all the ncRNA and gene nodes linked to kth gene. $\Sigma_{Y_k \mathcal{N}(Y_k)}$ is the vector of covariance between gene $Y_k$ and its directly linked nodes. $\Sigma_{Y_k}$ and $\Sigma_{\mathcal{N}(Y_k)}$ are the variance of $Y_k$ and the diagonal matrix of variances of its directly linked nodes. Nodes with small $\phi$'s will be removed as determined by Wilk's Lambda asymptotic test. Detailed algorithm can be found in Algorithm S4.

**Algorithm S4:** Node-wise screening

**Input:** ncRNA expression data $\boldsymbol{X} = (X_1, ..., X_{p_1})$ and gene expression data $\boldsymbol{Y} = (Y_1, ..., Y_{q_1})$, partially directed graph $\mathcal{G}_1 = \{\mathcal{V}_1, \mathcal{E}_1\}$; Active sets of ncRNA and gene nodes: $\mathcal{V}_1^X$ and $\mathcal{V}_1^Y$, pre-determined $\alpha$ from algorithm S1. Let $\mathcal{V}_2 = \mathcal{V}_1$

**for** *each ncRNA node* $X_j \in \mathcal{V}_1^X$ **do**

  Compute $\phi_j^X = \max_a \frac{\Sigma_{X_j \mathcal{N}(X_j)} a}{\sqrt{\Sigma_{X_j}} \sqrt{a^T \Sigma_{\mathcal{N}(X_j)} a}}$, where $\mathcal{N}(X_j) = \{Y_k : (X_j \to Y_k) \in \mathcal{E}_1\}$ ;

  **if** $WL(\phi_j^X) > \alpha$, *where* $WL(\cdot)$ *gives the p-value for the Wilks' Lambda asymptotic test* **then**

    remove $X_j$ from $\mathcal{V}_2$

**for** *each gene node* $Y_k \in \mathcal{V}_1^Y$ **do**

  Compute $\phi_k^Y = \max_b \frac{\Sigma_{Y_k \mathcal{N}(Y_k)} b}{\sqrt{\Sigma_{Y_k}} \sqrt{b^T \Sigma_{\mathcal{N}(Y_k)} b}}$, where $\mathcal{N}(Y_k) = \{X_j : (X_j \to Y_k) \in \mathcal{E}_1\}$ ;

  **if** $WL(\phi_k^Y) > \alpha$, *where* $WL(\cdot)$ *gives the p-value for the Wilks' Lambda asymptotic test* **then**

    remove $Y_k$ from $\mathcal{V}_2$

**Output:** $\mathcal{V}_2$: the active set for remaining nodes.

#### S1.5 BN structure learning stage II: score-based order MCMC algorithm

After stage I, the number of edges and nodes thus the search space for DAG is greatly reduced. In stage II, we proposed a score-based order MCMC algorithm (Friedman and Koller, 2003) adapted to the semi-bipartite graph and incorporated the prior knowledge on ncRNA-disease association and ncRNA-gene interaction information to search the causal structure that best represents the data and curated database.

Suppose we had a partially directed graph $\mathcal{G}_2 = \{\mathcal{V}_2, \mathcal{E}_2\}$ left after stage I, where $\mathcal{V}_2 = X_1, ..., X_{p_2}, Y_1, ..., Y_{q_2}$, $p_2$ and $q_2$ are the number of remaining ncRNAs and genes, $\mathcal{E}_2$ includes all remaining edges, including both directed (ncRNA-gene directed edges and some gene-gene directed edges determined in stage I) and undirected edges (gene-gene edges not yet determined in stage I). We know that $p_2 \ll p, q_2 \ll q$ and $|\mathcal{E}_2| \ll pq + q(q-1)/2$ after stage

I.

Following Friedman and Koller (2003), denote by $\prec$ the topological order of nodes in a graph and $P(\mathcal{G}|D) = \prod_{k \in \mathcal{V}_\mathcal{G}} s(Y_k, Pa_\mathcal{G}(Y_k)|D)$ the Bayesian Gaussian equivalence (BGe) score in our context, where $D$ is the data, $s$ is the score function that only depends on gene nodes (e.g. $Y_k$) and their parent set (e.g. $Pa_{G_2}(Y_k)$) including both ncRNA and gene nodes. In order space, each order receives a score equal to the sum of the scores of all DAGs compatible with this order: $R(\prec|D) = \Sigma_{\mathcal{G} \in \Gamma_\prec} P(\mathcal{G}|D) = \Sigma_{\mathcal{G} \in \Gamma_\prec} \prod_{k \in \mathcal{V}_\mathcal{G}} s(Y_k, Pa_\mathcal{G}(Y_k)|D)$. The order MCMC algorithm will sample over space of orderings and obtain the overall probability of a particular feature occurring. At each iteration $t$, we propose a move to change the order: (1) global swap, in which any two nodes will swap positions in the node ordering while keeping other fixed; (2) local transposition, in which only two adjacent nodes will swap positions; or (3) node relocation, in which we place a single node in any position of the current order and accept with probability $\rho = \min\{1, \frac{R(\prec'|D)}{R\left(\prec^{[t]}|D\right)}\}$. Assuming that the Markov chain has converged within $T$ steps, and we obtain a sample of DAGs $\mathcal{G}^{(1)}, \ldots, \mathcal{G}^{(T)}$. After excluding the first $t'$ burn-in steps, we can approximate the posterior probability by the sample average and identify the final graph with the highest posterior probability (Kuipers et al., 2022). When applying the algorithm to real data examples, suppose we have a set of validated disease associated ncRNAs $\Omega_n$ and a set of validated ncRNA-gene edges $\Omega_e$ from curated database and we incorporated them as prior knowledge into our algorithm. During MCMC sampling, we sampled with a relatively higher probability for edges belonging to $\Omega_e$ or edges that involved nodes belonging to $\Omega_n$ so the validated disease-associated ncRNA nodes and ncRNA-gene regulatory pairs were more likely to be sampled and remained in the final network. Detailed algorithm is summarized in Algorithm S5.

**Algorithm S5:** Stage II order MCMC algorithm

**Input** ncRNA expression data $\boldsymbol{X} = (X_1, ..., X_{p_2})$ and gene expression data $\boldsymbol{Y} = (Y_1, ..., Y_{q_2})$, partially directed graph $\mathcal{G}_2 = \{\mathcal{V}_2, \mathcal{E}_2\}$ left after stage I, validated disease associated ncRNA nodes $\Omega_n$ and ncRNA-gene edges $\Omega_e$ from curated database.

**Initialize** node ordering $\prec^{[0]}= (\prec_X, \prec_Y) = (X_{(1)}, ..., X_{(p_2)}, Y_{(1)}, ..., Y_{(q_2)})$ and $t \leftarrow 0$

**repeat**

Given the current order $\prec^{[t]}$, propose a new order $\prec'$ is from any of the following moves:

- local move: swapping **adjacent** ncRNA or gene nodes in $\prec^{[t]}$.
- global move: swapping two **random** ncRNA or gene nodes in $\prec^{[t]}$.
- node relocation: place a single node in any position of the current order $\prec^{[t]}$.

The new order will be accepted with probability $\rho = \min\left\{1, \frac{R(\prec'|D)}{R(\prec^{[t]}|D)}\right\}$;
If accepted, $\prec^{[t+1]}=\prec'$; otherwise, $\prec^{[t+1]}=\prec^{[t]}$.

Given the new order $\prec^{[t+1]}$, we will sample a DAG consistent with the order by sampling the parents of each node proportionally to the entries respecting the order in the score table. If $(j^\dagger, k^\dagger)$th edge $e_{j^\dagger k^\dagger} \in \Omega_e$, it will have twice the probability to be sampled than the other edges; if $j^\ddagger$th node $\in \Omega_n$, then $\forall_k e_{j^\ddagger k}$ will have twice the probability to be sampled than the other edges.

$t \leftarrow t + 1$

**until** *convergence*;

**Output** A Markov chain of T samples of graphs $\mathcal{G}^{(1)}, \mathcal{G}^{(2)}, \ldots, \mathcal{G}^{(T)}$.

We will then take the sample average to approximate the posterior probability and identify the final graph with the highest posterior probability.

#### S1.6 Differential ncRNA regulatory network analysis

The ncRNA regulatory network could vary condition by condition implying the difference in underlying molecular regulatory mechanisms. Running our algorithm independently in two conditions generates two completely different regulatory network structures that are hardly comparable and direct search of a global differential regulatory network is computationally infeasible (Grimes et al., 2019). In this paper, we borrowed the incremental learning idea (Van de Ven et al., 2022): we first used one condition as reference to generate an initial regulatory network by applying the two-stage algorithm; then we updated the network using the data from the other condition by applying the proposed MCMC sampling scheme; this new network will be compared to the initial network for difference in connection patterns (e.g. what edges gained and lost).

**Algorithm S6:** Identifying differential regulatory network between two conditions

**Input:** ncRNA and gene expression data from condition 1 and 2, denoted as $D_{cond1}$ with sample size $n_1$ and $D_{cond2}$ with sample size $n_2$, respectively.

**Step 1. Generate a reference network from condition 1.**

Run Algorithm S5 on the data from condition 1 $D_{cond1}$ to generate a reference network for condition 1 $\mathcal{G}_{cond1}$. As a byproduct, obtain the ordering of nodes at the convergence $\prec_{cond1}$.

**Step 2. Update the reference network using data from condition 2.**

Suppose $\prec'$ is the proposed new order, we will run Algorithm S5 again now using data from condition 2 $D_{cond2}$ and initial ordering $\prec_{cond1}$. The new order will have the acceptance probability:

$$\rho = \min\left\{1, \frac{R\left(\prec' \mid D_{cond2}\right)}{R\left(\prec_{cond1} \mid D_{cond2}\right)}\right\}$$

This way we will have an updated network $\mathcal{G}_{cond2}$ based on new data in $D_{cond2}$.

**Step 3. Subsampling to mitigate bias from imbalanced sample size.**

Suppose the reference condition 1 has larger sample size, i.e. $n_1 > n_2$, take $B = 100$ subsamples of size $n_2$ from $D_{cond1}$, denoted as $D^{(1)}_{cond1}, ..., D^{(B)}_{cond1}$.

**for** *each* $D^{(b)}_{cond1}$ **do**

  Run Step 2 and obtain a network $\mathcal{G}^{(b)}_{cond1}$ .

**for** *each edge* $j \to k$ **do**

  calculate

  $\phi(j \to k) = \frac{1}{B}\sum_{b=1}^{B} \mathbf{1}\{(j \to k)$ appears in exact one of $\mathcal{G}_{cond1}$ and $\mathcal{G}^{(b)}_{cond1}\}$.

We define $(j \to k)$ edge as differential edge if $\phi(j \to k) < 0.5$ and $(j \to k)$ appears in exact one of $\mathcal{G}_{cond1}$ and $\mathcal{G}_{cond2}$.

#### S1.7 Connection between partial correlation, multivariate regression and semi-bipartite graph

In our first stage of BN structure learning, we utilize a robust version of partial correlation used to screen for predictor-response pairs in multivariate regression to screen for edges in semi-bipartite graph. In this section, we will briefly discuss the connection between partial correlation, multivariate regression and semi-bipartite graph.

For a multivariate regression model:

$$Y_{n\times q} = X_{n\times p}\beta_{p\times q} + E_{n\times q}, E \sim N(0, \Sigma_Y \otimes I_n),$$

where $\beta$ is the $p \times q$ matrix of regression coefficients, $cov(X) = \Sigma_X$, $\Sigma_Y$ is the $q \times q$ covariance matrix of $Y$. Suppose $T = (X\ Y)$ and $cov(T) = \Sigma$, with block-wise decomposition, $\Sigma = \begin{pmatrix} \Sigma_X & \Sigma_{XY} \\ \Sigma_{YX} & \Sigma_Y \end{pmatrix}$. Assuming $\Sigma$ is strictly positive definite thus invertible, and let $\Theta = \Sigma^{-1} = \begin{pmatrix} \Theta_X & \Theta_{XY} \\ \Theta_{YX} & \Theta_Y \end{pmatrix}$. It can be shown that the above multivariate regression model can be reparameterized in the form of a conditional Gaussian Graphical Model (cGGM) (Sohn and Kim, 2012; Chiquet et al., 2017):

$$Y|X \sim N(-X\Theta_{XY}\Theta_Y^{-1}, \Theta_Y^{-1} \otimes I_n),$$

where $\beta = -\Theta_{XY}\Theta_Y^{-1}$. The partial correlation $\rho(X_j, Y_k|X^{\{j\}^C}, Y^{\{k\}^C}) = -\frac{\Theta_{X_jY_k}}{\Theta_{X_jX_j}\Theta_{Y_kY_k}}$, so we have $\rho(X_j, Y_k|X^{\{j\}^C}, Y^{\{k\}^C}) = 0 \leftrightarrow \Theta_{X_jY_k} = 0$, thus there is a one-to-one connection between the regression coefficient in multivariate regression and the full partial correlation which can be represented by precision in cGGM.

Semi-bipartite graph models the connection between $X$ and $Y$ nodes as reflected in $\beta$ in cGGM. In the same time, in our semi-bipartite graph, the set of $Y$ nodes is allowed to be connected with each other, so the internal dependencies among $Y$ nodes can be explicitly modeled by $\Sigma_Y$ as shown above. So in practice, cGGM can be used to estimate edges in a semi-bipartite graph based on normally distributed data (Gross and Sullivant, 2018).

### S2 Supplementary: simulation

#### S2.1 Simulation setting

In this section, we conducted simulation studies to benchmark the performance of our method compared to other popular gene regulatory network detection methods including correlation-based methods (Pearson), machine learning based methods (GENIE3 (Huynh-Thu et al., 2010)) and other Bayesian network methods (constraint-based PC (PC) algorithm (Kalisch and Bühlman, 2007) and Max-Min Parents Children (MMPC) algorithm (Tsamardinos et al., 2003), score-based Hill-Climbing (HC, (Heckerman et al., 1995)), and hybrid approaches Hybrid MCMC (Kuipers et al., 2022) and Adaptively Restricted Greedy Equivalence Search (ARGES (Nandy et al., 2018)). Following the notation from main text, denote the ncRNA expression data by $\boldsymbol{X_{n\times p}} = (X_1, ..., X_p)$ and the gene expression data as $\boldsymbol{Y_{n\times q}} = (Y_1, ..., Y_q)$ where $p$ and $q$ are numbers of ncRNAs and genes, respectively, and $n$ is the sample size. A DAG can be represented with an adjacency matrix:

$$\boldsymbol{A}_{(p+q)\times(p+q)} = (A_{jk}) = \begin{pmatrix} \boldsymbol{A}^{(XX)} & \boldsymbol{A}^{(XY)} \\ \boldsymbol{A}^{(YX)} & \boldsymbol{A}^{(YY)} \end{pmatrix},$$

where $A_{jk} = 1$ indicates there is an edge going from j-th node to k-th node. The adjacency matrix $\boldsymbol{A}_{(p+q)\times(p+q)}$ can be partitioned into four parts, each of which represents one type of edge in ncRNA-gene regulatory networks. Given the semi-bipartite graph structure of ncRNA regulatory network we assumed here, we will have $\boldsymbol{A}^{(XX)} = \boldsymbol{A}^{(YX)} = \boldsymbol{0}$, i.e. no ncRNAs are regulated by genes and we do not consider the direct interaction between two ncRNAs. The remaining two types of edges in our studies include:

- Level-1: edges from ncRNAs to genes, denoted in $\boldsymbol{A}^{(XY)}$
- Level-2: edges from genes to genes, denoted in $\boldsymbol{A}^{(YY)}$

We first simulated the ncRNA expression data from normal distribution with mean 0 and variance 0.2. To generate gene expression data, we simulated the adjacency matrix according to a repetitive pattern as shown in Figure S3. For level-1 edges, we assumed there existed "hub ncRNAs" serving as master regulators that regulate multiple genes simultaneously, commonly observed in existing studies (Statello et al., 2021; Plaisier et al., 2012). We then randomly picked gene → gene regulation relationships and generated level-2 edges. The gene expression data can be generated based on

$$Y_k = \sum_{j=1}^{p} \boldsymbol{A}_{jk}^{(XY)} X_j + \sum_{k'\neq k} \boldsymbol{A}_{k'k}^{(YY)} Y_{k'} + \varepsilon \text{ and } \varepsilon \sim N(0, 0.2), \text{ for } k = 1, ..., q.$$

Two scenarios with different dimensions are considered:

- Low-dimension: $n = 200$, $p = q = 100$, number of true edges is 100.
- High-dimension: $n = 200$, $p = q = 1000$, number of true edges is 100.

The DAGs identified by each method were evaluated based on sensitivity/recall (True Positive Rate) and precision (Positive Predictive Value) as well as the F-score:

$$\text{sensitivity} = \frac{TP}{P}, \text{precision} = \frac{TP}{TP + FP}.$$

Note that when calculating sensitivity and precision, we consider our DAGs/adjacency matrices to be directed here. Specifically, $TP$ indicates edges in the true network that are also identified by the method and with the correct direction. Conversely, $FP$ indicates either edges that are not in the true network but identified by the method, or edges in the true network but identified by the method in the wrong direction.

In addition, F-scores were also computed as an overall evaluation of how well the structure learning algorithms balance between capturing true regulatory relationship (sensitivity) and forcing sparse DAGs (precision):

$$F - score = \frac{2 * precision * sensitivity}{precision + sensitivity}.$$

Each method has a different cutoff, we hereby drew the plots over a range of thresholds according to different tuning parameters for each method for a fair comparison. Each simulation scenario is repeated for 100 times and we compared the average results.

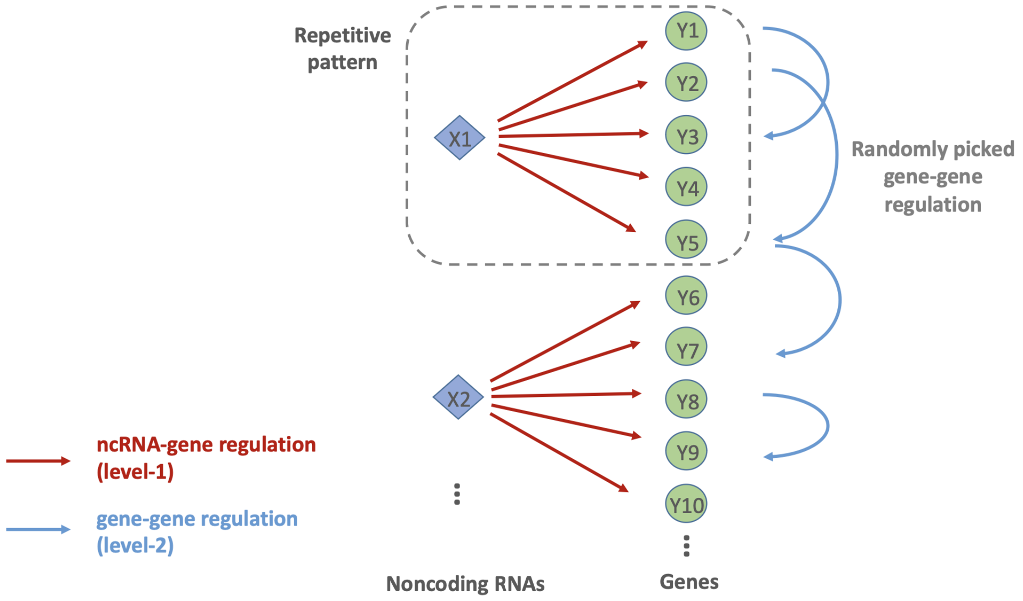


Figure S3: Repetitive pattern that was used to generate DAGs in simulation studies.

#### S2.2 Simulation results

Simulation results were presented in Figure S4 and S5. In the low-dimension scenario, our CAR-NET method better recovered the true DAG with high sensitivity while maintaining graph sparsity (relatively high precision) similar to other BN methods such as HC, PC and Hybrid MCMC (Figure S4 left). CAR-NET method also had the highest F score over almost all choices of tuning parameter (especially for level-1 and all edges), indicating the great overall performance of our method (Figure S4 right). Though Pearson and GENIE3 methods can sometimes reach high sensitivity, overall they did poorly in recovering the true network with low F scores and achieved high sensitivity at the cost of low precision resulting in a large dense network harder to interpret. Our method was particularly advantageous in level-1 edge selection (Figure S4 (b)), as our method specifically incorporated the semi-bipartite structure of the ncRNA-gene regulation problem into our network discovery algorithm. In the high-dimensional scenario, except for Pearson method and the two constraint-based BN methods PC and MMPC, other methods ran into computational bottleneck thus were not included for evaluation. The advantage of CAR-NET has become more clear in the high-dimension setting with both a higher sensitivity and precision when compared to other methods. The low computational cost of our two-stage method also makes it attractive to handle high-dimensional data with wider applicability.

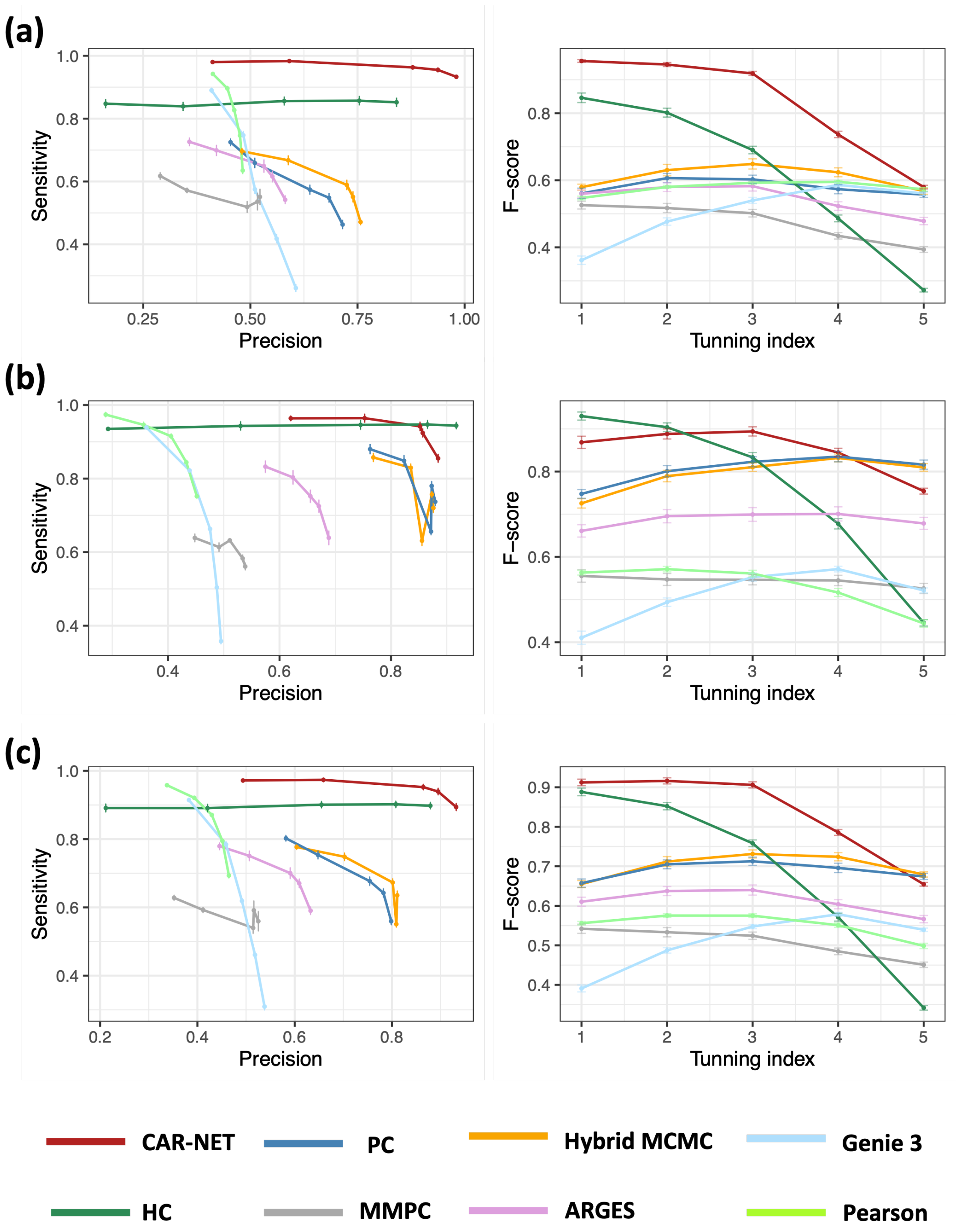


Figure S4: Simulation results in low-dimension setting for: (a) level-1 edges; (b) level-2 edges; (c) all edges. The left panel is sensitivity vs. precision plot and the right panel is F-score comparison.

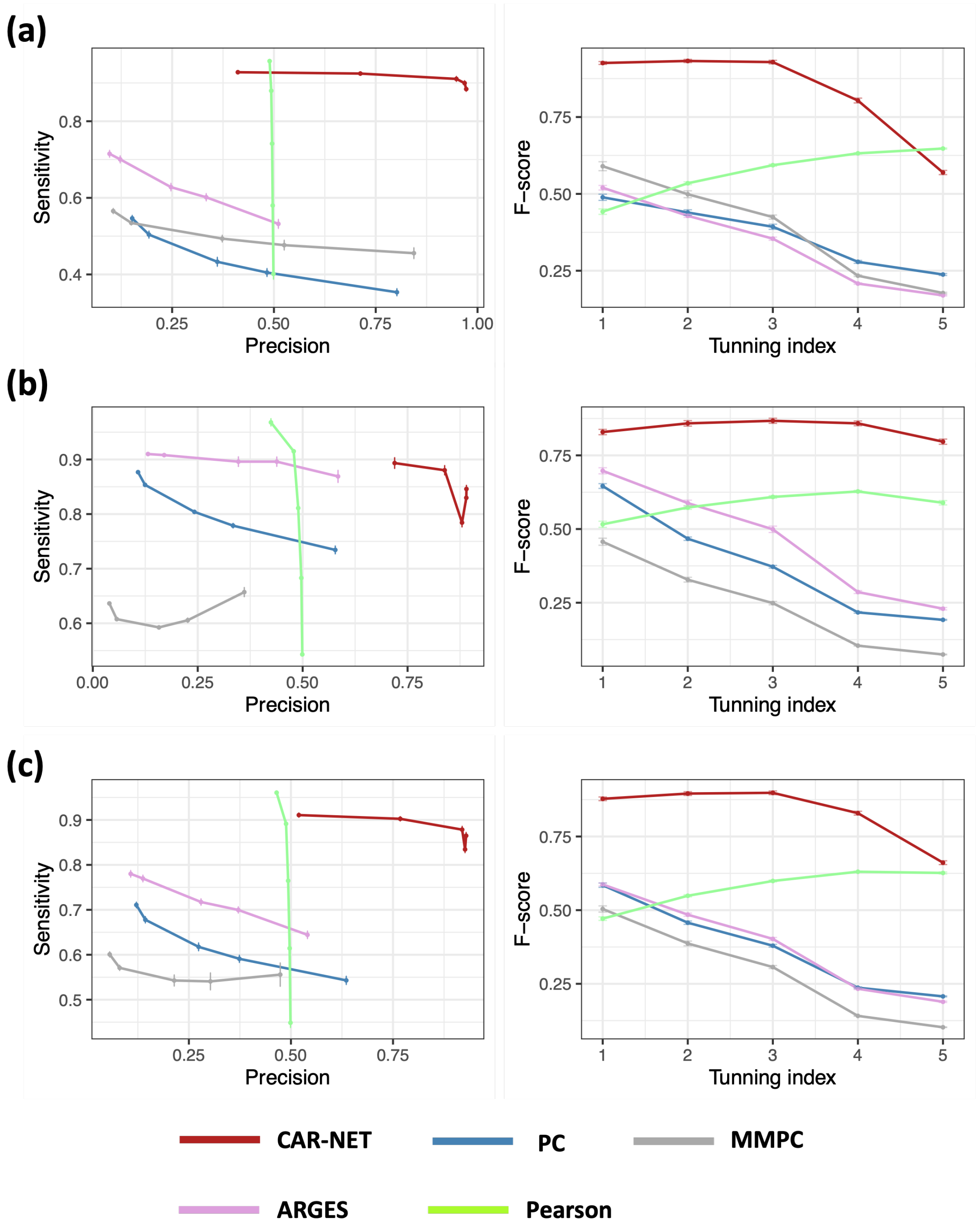


Figure S5: Simulation results in high-dimension setting for: (a) level-1 edges; (b) level-2 edges; (c) all edges. The left panel is sensitivity vs. precision plot and the right panel is F-score comparison.

#### References


Bühlmann, P., Kalisch, M., and Maathuis, M. H. (2010). Variable selection in high-dimensional linear models: partially faithful distributions and the pc-simple algorithm. *Biometrika*, 97(2):261–278.

Chickering, D. M. (2002). Learning equivalence classes of bayesian-network structures. *Journal of machine learning research*, 2(Feb):445–498.

Chiquet, J., Mary-Huard, T., and Robin, S. (2017). Structured regularization for conditional gaussian graphical models. *Statistics and Computing*, 27(3):789–804.

Cohen, J. (2013). *Statistical power analysis for the behavioral sciences*. routledge.

Friedman, N. and Koller, D. (2003). Being bayesian about network structure. a bayesian approach to structure discovery in bayesian networks. *Machine learning*, 50:95–125.

Grimes, T., Potter, S. S., and Datta, S. (2019). Integrating gene regulatory pathways into differential network analysis of gene expression data. *Scientific reports*, 9(1):5479.

Gross, E. and Sullivant, S. (2018). The maximum likelihood threshold of a graph.

Hauser, A. and Bühlmann, P. (2012). Characterization and greedy learning of interventional markov equivalence classes of directed acyclic graphs. *The Journal of Machine Learning Research*, 13(1):2409–2464.

Heckerman, D., Geiger, D., and Chickering, D. M. (1995). Learning bayesian networks: The combination of knowledge and statistical data. *Machine learning*, 20:197–243.

Huynh-Thu, V. A., Irrthum, A., Wehenkel, L., and Geurts, P. (2010). Inferring regulatory networks from expression data using tree-based methods. *PloS one*, 5(9):e12776.

Kalisch, M. and Bühlman, P. (2007). Estimating high-dimensional directed acyclic graphs with the pc-algorithm. *Journal of Machine Learning Research*, 8(3).

Ke, H., Ren, Z., Qi, J., Chen, S., Tseng, G. C., Ye, Z., and Ma, T. (2022). High-dimension to high-dimension screening for detecting genome-wide epigenetic and noncoding rna regulators of gene expression. *Bioinformatics*, 38(17):4078–4087.

Koller, D. and Friedman, N. (2009). *Probabilistic graphical models: principles and techniques*. MIT press.

Kuipers, J., Suter, P., and Moffa, G. (2022). Efficient sampling and structure learning of bayesian networks. *Journal of Computational and Graphical Statistics*, 31(3):639–650.

Ma, T., Ke, H., and Ren, Z. (2022). Robust distance correlation for variable screening. *arXiv preprint arXiv:2212.13292*.

Nandy, P., Hauser, A., and Maathuis, M. H. (2018). High-dimensional consistency in score-based and hybrid structure learning. *The Annals of Statistics*, 46(6A):3151–3183.

Pearl, J. (2009). *Causality*. Cambridge university press.

Pearl, J. (2014). *Probabilistic reasoning in intelligent systems: networks of plausible inference*. Elsevier.

Plaisier, C. L., Pan, M., and Baliga, N. S. (2012). A mirna-regulatory network explains how dysregulated mirnas perturb oncogenic processes across diverse cancers. *Genome research*, 22(11):2302–2314.

Sohn, K.-A. and Kim, S. (2012). Joint estimation of structured sparsity and output structure in multiple-output regression via inverse-covariance regularization. In *Artificial Intelligence and Statistics*, pages 1081–1089. PMLR.

Spirtes, P., Glymour, C., and Scheines, R. (2001). *Causation, prediction, and search*. MIT press.

Spirtes, P. L., Meek, C., and Richardson, T. S. (2013). Causal inference in the presence of latent variables and selection bias. *arXiv preprint arXiv:1302.4983*.

Statello, L., Guo, C.-J., Chen, L.-L., and Huarte, M. (2021). Gene regulation by long non-coding rnas and its biological functions. *Nature reviews Molecular cell biology*, 22(2):96–118.

Tsamardinos, I., Aliferis, C. F., and Statnikov, A. (2003). Time and sample efficient discovery of markov blankets and direct causal relations. In *Proceedings of the ninth ACM SIGKDD international conference on Knowledge discovery and data mining*, pages 673–678.

Tsamardinos, I., Brown, L. E., and Aliferis, C. F. (2006). The max-min hill-climbing bayesian network structure learning algorithm. *Machine learning*, 65:31–78.

Van de Ven, G. M., Tuytelaars, T., and Tolias, A. S. (2022). Three types of incremental learning. *Nature Machine Intelligence*, 4(12):1185–1197.

Veličković, P., Cucurull, G., Casanova, A., Romero, A., Lio, P., and Bengio, Y. (2017). Graph attention networks. *arXiv preprint arXiv:1710.10903*.

Vogels, L., Mohammadi, R., Schoonhoven, M., and Birbil, Ş. İ. (2024). Bayesian structure learning in undirected gaussian graphical models: Literature review with empirical comparison. *Journal of the American Statistical Association*, pages 1–19.
