## Supplementary tables for "Inferring non-coding RNA regulatory network from transcriptomic data and curated databases"

| Table S1. Summary list of curated ncRNA databases | | | |
|---|---|---|---|
| **Database** | **ncRNA type** | **Description** | **Link** |
| miRTarBase | miRNA | The experimentally validated microRNA-target interactions database | https://mirtarbase.cuhk.edu.cn/ |
| miRecords | miRNA | A large, high-quality manually curated database of experimentally validated miRNA-target interactions | http://mirecords.umn.edu/miRecords |
| TargetScan | miRNA | A database of miRNA target predictions across species | https://www.targetscan.org/vert_80/ |
| miRCancer | miRNA | Comprehensive collection of miRNA associations with various human cancers automatically extracted from published literatures in PubMed. | http://mircancer.ecu.edu/ |
| miR2Disease | miRNA | A manually curated database that provides a comprehensive resource of miRNA deregulation in various human diseases | http://www.mir2disease.org/ |
| LNCipedia | lncRNA | Comprehensive database for human lncRNAs, including annotations and structure. | https://lncipedia.org/ |
| EVLncRNAs | lncRNA | A comprehensive and manually curated resource of integrated sequence, structure, functional, and phenotypic information of experimentally validated lncRNAs | https://www.sdklab-biophysics-dzu.net/EVLncRNAs3/#/ |
| LncRNA2Target | lncRNA | A comprehensive database for target genes of lncRNAs in human and mouse. It collects lncRNA-target relationships from papers and lncRNA knockdown/overexpression RNA-seq datasets. | http://123.59.132.21/lncrna2target |

| | | | |
|---|---|---|---|
| LncRNADisease | lncRNA, circRNA | A database of comprehensive experimentally supported and predicted ncRNA-disease associations curated from manual literatures and other resources | http://www.rnanut.net/lncrnadisease/ |
| LncTarD | lncRNA | A database for experimentally supported key lncRNA-target regulations, their influenced functions and lncRNA-mediated regulatory mechanisms in human diseases | https://lnctard.bio-database.com/ |
| LncBook | lncRNA | A human lncRNA database with functional and disease-related information. | https://ngdc.cncb.ac.cn/lncbook/ |
| snoDB | snoRNA | An interactive database of human snoRNAs that includes up-to-date information on snoRNA features, genomic location, conservation, host gene, snoRNA–RNA targets and snoRNA abundance | https://bioinfo-scottgroup.med.usherbrooke.ca/snoDB/ |
| snoRNA-LBME-db | snoRNA | A dedicated database containing human C/D box and H/ACA box small nucleolar RNAs (snoRNAs), and small Cajal body-specific RNAs (scaRNAs) | http://www-snorna.biotoul.fr/ |
| sRNAMap | snRNA | Genomic maps for small non-coding RNAs, their regulators and their targets in microbial genomes | http://sRNAMap.mbc.nctu.edu.tw/ |
| Circ2Disease | circRNA | A manually curated database of experimentally validated circRNAs in human disease | http://cgga.org.cn:9091/circRNADisease/ |
| CircR2Cancer | circRNA | A manually curated database of associations between circRNAs and cancers | https://ngdc.cncb.ac.cn/databasecommons/database/id/7942 |

**Table S2. Summary of Case Study 1**

| Methods | lncRNA node | Gene node | Level-1 edge | Level-2 edge |
|---|---|---|---|---|
| CAR-NET | 408 | 588 | 669 | 509 |
| PC | 1,205 | 2,531 | 306 | 2,027 |
| GENIE3 | 3,031 | 4,511 | 75,020 | 342,923 |
| WGCNA | 3,367 | 4,511 | 2,228,021 | 4,000,270 |
| Pearson | 3,367 | 4,511 | 8,728,171 | 13,052,722 |

### Table S3. Pathway analysis results for Case Study 1 (BH adjusted p-value < 0.05)

| pathway | pvalue | BH adjusted pvalue | pathway_size | # genes |
|---|---|---|---|---|
| GO:CC endosome | 0.00341 | 0.03882 | 21 | 8 |
| Reactome Sema4D in semaphorin signaling | 0.0057 | 0.03882 | 14 | 6 |
| Reactome RNA Polymerase III Transcription | 0.00871 | 0.03882 | 11 | 5 |
| Reactome GPVI-mediated activation cascade | 0.00871 | 0.03882 | 11 | 5 |
| GO:CC actin cytoskeleton | 0.00982 | 0.03882 | 50 | 13 |
| Reactome RNA Polymerase III Transcription Termination | 0.01299 | 0.03882 | 8 | 4 |
| GO:CC dendrite | 0.01335 | 0.03882 | 12 | 5 |
| GO:BP regulation of myeloid cell differentiation | 0.018 | 0.03882 | 5 | 3 |
| GO:CC late endosome | 0.018 | 0.03882 | 5 | 3 |
| Reactome Membrane binding and targetting of GAG proteins | 0.018 | 0.03882 | 5 | 3 |
| Reactome p75NTR signals via NF-kB | 0.018 | 0.03882 | 5 | 3 |
| Reactome Sema4D induced cell migration and growth-cone collapse | 0.01941 | 0.03882 | 13 | 5 |
| Reactome Interleukin-3; 5 and GM-CSF signaling | 0.01941 | 0.03882 | 13 | 5 |
| BioCarta Role of PI3K subunit p85 in regulation of Actin Organization | 0.01941 | 0.03882 | 13 | 5 |
| GO:MF sequence-specific DNA binding transcription factor activity | 0.02095 | 0.03911 | 90 | 19 |
| BioCarta Y branching of actin filaments | 0.02704 | 0.04143 | 14 | 5 |
| GO:BP amino acid transport | 0.03147 | 0.04143 | 10 | 4 |
| GO:BP myeloid cell differentiation | 0.03147 | 0.04143 | 10 | 4 |
| Reactome TGF-beta receptor signaling in EMT (epithelial to mesench | 0.03255 | 0.04143 | 6 | 3 |
| Reactome Nuclear Receptor transcription pathway | 0.03255 | 0.04143 | 6 | 3 |
| Reactome CD28 dependent Vav1 pathway | 0.03255 | 0.04143 | 6 | 3 |
| Reactome Regulation of signaling by CBL | 0.03255 | 0.04143 | 6 | 3 |
| KEGG Phosphatidylinositol signaling system | 0.036 | 0.04237 | 25 | 7 |
| Reactome G1 Phase | 0.03632 | 0.04237 | 15 | 5 |
| Reactome SMAD2/SMAD3:SMAD4 heterotrimer regulates transcriptic | 0.04447 | 0.04612 | 11 | 4 |
| Reactome CD28 co-stimulation | 0.04447 | 0.04612 | 11 | 4 |
| BioCarta How does salmonella hijack a cell | 0.04447 | 0.04612 | 11 | 4 |
| BioCarta Influence of Ras and Rho proteins on G1 to S Transition | 0.04734 | 0.04734 | 16 | 5 |

### Table S4. Summary of Case Study 2

| | | | | |
|---|---|---|---|---|
| Final network for early-stage | | | | |
| **Methods** | **miRNA node** | **Gene node** | **Level-1 edge** | **Level-2 edge** |
| CAR-NET | 127 | 377 | 328 | 260 |
| PC | 382 | 3,635 | 61 | 3417 |
| GENIE3 | 738 | 7,311 | 104,297 | 4,415,376 |
| WGCNA | 750 | 7,311 | 882,671 | 6,600,966 |
| Pearson | 750 | 7,311 | 1,211,724 | 24,625,122 |
| Differential network analysis results from CAR-NET | | | | |
| | **miRNA node** | **Gene node** | **Level-1 edge** | **Level-2 edge** |
| Early stage | 127 | 377 | 328 | 260 |
| Late stage | 89 | 264 | 153 | 116 |

**Table S5. Pathway analysis results for Case Study 2 (BH adjusted p-value < 0.05)**

| pathway | pvalue | BH adjusted pvalue | pathway_size | # genes |
|---|---|---|---|---|
| KEGG Ubiquitin mediated proteolysis | 0.00172 | 0.0462 | 26 | 6 |
| KEGG Amino sugar and nucleotide sugar metabolism | 0.00247 | 0.0462 | 12 | 4 |
| Reactome Platelet calcium homeostasis | 0.00408 | 0.0462 | 7 | 3 |
| KEGG Arginine and proline metabolism | 0.00612 | 0.0462 | 33 | 6 |
| GO:MF small protein conjugating enzyme activity | 0.00905 | 0.0462 | 9 | 3 |
| GO:MF ubiquitin-like protein transferase activity | 0.00905 | 0.0462 | 9 | 3 |
| GO:MF ubiquitin-protein transferase activity | 0.00905 | 0.0462 | 9 | 3 |
| GO:BP response to nutrient | 0.01244 | 0.0462 | 10 | 3 |
| GO:MF acid-amino acid ligase activity | 0.01244 | 0.0462 | 10 | 3 |
| BioCarta Adhesion and Diapedesis of Granulocytes | 0.01647 | 0.0462 | 11 | 3 |
| GO:MF enzyme inhibitor activity | 0.01652 | 0.0462 | 52 | 7 |
| GO:MF sugar binding | 0.01747 | 0.0462 | 20 | 4 |
| GO:MF ligase activity | 0.02073 | 0.0462 | 21 | 4 |
| GO:MF ligase activity; forming carbon-nitrogen bonds | 0.02113 | 0.0462 | 12 | 3 |
| GO:BP photoreceptor cell maintenance | 0.02391 | 0.0462 | 5 | 2 |
| GO:BP regulation of protein stability | 0.02391 | 0.0462 | 5 | 2 |
| KEGG RNA degradation | 0.02391 | 0.0462 | 5 | 2 |
| BioCarta Role of Parkin in the Ubiquitin-Proteasomal Pathway | 0.02391 | 0.0462 | 5 | 2 |
| GO:BP response to nutrient levels | 0.02644 | 0.0462 | 13 | 3 |
| GO:BP cell recognition | 0.02644 | 0.0462 | 13 | 3 |
| GO:MF hydro-lyase activity | 0.02644 | 0.0462 | 13 | 3 |
| GO:BP sensory perception | 0.02858 | 0.0462 | 58 | 7 |
| GO:BP G-protein coupled receptor signaling pathway; coupled to cyclic nucle | 0.02878 | 0.0462 | 34 | 5 |
| GO:BP cyclic-nucleotide-mediated signaling | 0.03221 | 0.0462 | 35 | 5 |
| GO:MF carbon-oxygen lyase activity | 0.03239 | 0.0462 | 14 | 3 |
| Reactome Smooth Muscle Contraction | 0.03239 | 0.0462 | 14 | 3 |
| GO:BP insulin receptor signaling pathway | 0.03465 | 0.0462 | 6 | 2 |
| GO:CC dystrophin-associated glycoprotein complex | 0.03465 | 0.0462 | 6 | 2 |
| GO:MF insulin-like growth factor receptor binding | 0.03465 | 0.0462 | 6 | 2 |
| Reactome Ethanol oxidation | 0.03465 | 0.0462 | 6 | 2 |
| Reactome Platelet homeostasis | 0.0398 | 0.04808 | 37 | 5 |
| GO:BP embryo development | 0.04234 | 0.04808 | 26 | 4 |
| GO:BP response to extracellular stimulus | 0.0462 | 0.04808 | 16 | 3 |

| | | | | |
|---|---|---|---|---|
| GO:BP mesoderm development | 0.0462 | 0.04808 | 16 | 3 |
| GO:BP peripheral nervous system development | 0.04688 | 0.04808 | 7 | 2 |
| GO:CC microvillus | 0.04688 | 0.04808 | 7 | 2 |
| GO:MF oxidoreductase activity; acting on the CH-NH group of donors | 0.04688 | 0.04808 | 7 | 2 |
| GO:MF carbonate dehydratase activity | 0.04688 | 0.04808 | 7 | 2 |
| Reactome P2Y receptors | 0.04688 | 0.04808 | 7 | 2 |
| Reactome PPARA Activates Gene Expression | 0.04839 | 0.04839 | 39 | 5 |

### Table S6. Summary of Case Study 3

| Cell lines | ncRNA node | Gene node | Level-1 edge | Level-2 edge |
|---|---|---|---|---|
| Fibroblasts | 175 | 218 | 295 | 57 |
| HEK293T | 143 | 202 | 250 | 55 |
| MCF7 | 82 | 252 | 242 | 96 |

**Table S7. Pathway analysis results for fibroblast cell line in Case Study 3 (BH adjusted p-value < 0.05)**

| pathway | pvalue | BH adjusted pvalue | pathway_size | # genes |
|---|---|---|---|---|
| BioCarta Presenilin action in Notch and Wnt signaling | 0.00311 | 0.01736 | 5 | 3 |
| KEGG RIG-I-like receptor signaling pathway | 0.00359 | 0.01736 | 10 | 4 |
| GO:MF NF-kappaB binding | 0.00589 | 0.01736 | 6 | 3 |
| Reactome TRAF6 mediated IRF7 activation | 0.00589 | 0.01736 | 6 | 3 |
| Reactome TRAF6 mediated NF-kB activation | 0.00589 | 0.01736 | 6 | 3 |
| Reactome TRAF3-dependent IRF activation pathway | 0.00589 | 0.01736 | 6 | 3 |
| BioCarta ALK in cardiac myocytes | 0.00589 | 0.01736 | 6 | 3 |
| GO:BP regulation of kinase activity | 0.00651 | 0.01736 | 25 | 6 |
| GO:BP regulation of protein kinase activity | 0.00651 | 0.01736 | 25 | 6 |
| GO:BP regulation of transferase activity | 0.00797 | 0.01913 | 26 | 6 |
| GO:BP carbohydrate metabolic process | 0.00966 | 0.02108 | 27 | 6 |
| GO:BP cellular carbohydrate metabolic process | 0.01074 | 0.02148 | 20 | 5 |
| KEGG Cytosolic DNA-sensing pathway | 0.01483 | 0.02738 | 8 | 3 |
| GO:BP one-carbon metabolic process | 0.01766 | 0.02826 | 15 | 4 |
| GO:BP MAPK cascade | 0.01766 | 0.02826 | 15 | 4 |
| GO:BP protein kinase cascade | 0.01949 | 0.02924 | 49 | 8 |
| GO:BP positive regulation of transferase activity | 0.02862 | 0.0404 | 10 | 3 |
| Reactome RIG-I/MDA5 mediated induction of IFN-alpha/beta pathway | 0.03354 | 0.04472 | 18 | 4 |
| Reactome Negative regulators of RIG-I/MDA5 signaling | 0.03735 | 0.04478 | 11 | 3 |
| GO:BP regulation of catabolic process | 0.04291 | 0.04478 | 5 | 2 |
| GO:BP ER-nucleus signaling pathway | 0.04291 | 0.04478 | 5 | 2 |
| KEGG Starch and sucrose metabolism | 0.04291 | 0.04478 | 5 | 2 |
| Reactome Resolution of AP sites via the single-nucleotide replacement | 0.04291 | 0.04478 | 5 | 2 |
| GO:BP regulation of gene expression; epigenetic | 0.04727 | 0.04727 | 12 | 3 |

| Table S8. Pathway analysis results for HEK293 cell line in Case Study 3 (BH adjusted p-value < 0.05) | | | | |
|---|---|---|---|---|
| **pathway** | **pvalue** | **BH adjusted pvalue** | **pathway_size** | **# genes** |
| GO:MF oxidoreductase activity; acting on the CH-OH group of donors; NAD or NADP a | 0.00481 | 0.03124 | 18 | 5 |
| GO:MF oxidoreductase activity; acting on CH-OH group of donors | 0.00618 | 0.03124 | 19 | 5 |
| GO:BP cellular carbohydrate metabolic process | 0.00781 | 0.03124 | 20 | 5 |
| Reactome Triglyceride Biosynthesis | 0.02339 | 0.04466 | 10 | 3 |
| GO:BP carbohydrate metabolic process | 0.02805 | 0.04466 | 27 | 5 |
| GO:MF nucleotide kinase activity | 0.03722 | 0.04466 | 5 | 2 |
| GO:MF steroid binding | 0.03722 | 0.04466 | 5 | 2 |
| GO:MF organic acid transmembrane transporter activity | 0.03722 | 0.04466 | 5 | 2 |
| GO:MF carboxylic acid transmembrane transporter activity | 0.03722 | 0.04466 | 5 | 2 |
| BioCarta Oxidative Stress Induced Gene Expression Via Nrf2 | 0.03722 | 0.04466 | 5 | 2 |
| Reactome Signaling by Rho GTPases | 0.04395 | 0.04795 | 21 | 4 |
| KEGG Amino sugar and nucleotide sugar metabolism | 0.04824 | 0.04824 | 13 | 3 |

**Table S9. Pathway analysis results for MCF7 cell line in Case Study 3 (BH adjusted p-value < 0.05)**

| pathway | pvalue | BH adjusted pvalue | pathway_size | # genes |
|---|---|---|---|---|
| KEGG Cell cycle | 0.00795 | 0.04362 | 52 | 10 |
| Reactome Amino acid transport across the plasma membrane | 0.00888 | 0.04362 | 6 | 3 |
| Reactome Amino acid and oligopeptide SLC transporters | 0.00888 | 0.04362 | 6 | 3 |
| Reactome Mitotic G2-G2/M phases | 0.00987 | 0.04362 | 38 | 8 |
| Reactome Loss of Nlp from mitotic centrosomes | 0.01256 | 0.04362 | 32 | 7 |
| GO:CC microtubule organizing center | 0.01301 | 0.04362 | 25 | 6 |
| Reactome Transport of inorganic cations/anions and amino acids/oligopeptides | 0.01461 | 0.04362 | 7 | 3 |
| GO:BP organ development | 0.017 | 0.04362 | 58 | 10 |
| Reactome Recruitment of mitotic centrosome proteins and complexes | 0.01746 | 0.04362 | 34 | 7 |
| KEGG Porphyrin and chlorophyll metabolism | 0.02199 | 0.04362 | 8 | 3 |
| KEGG B cell receptor signaling pathway | 0.0224 | 0.04362 | 14 | 4 |
| GO:BP system development | 0.02624 | 0.04362 | 89 | 13 |
| GO:CC centrosome | 0.02874 | 0.04362 | 22 | 5 |
| Reactome ARMS-mediated activation | 0.03104 | 0.04362 | 9 | 3 |
| GO:CC Golgi apparatus | 0.03145 | 0.04362 | 55 | 9 |
| KEGG Axon guidance | 0.03581 | 0.04362 | 16 | 4 |
| GO:BP microtubule-based process | 0.03587 | 0.04362 | 31 | 6 |
| GO:CC DNA-directed RNA polymerase II; holoenzyme | 0.04062 | 0.04362 | 24 | 5 |
| KEGG Thyroid cancer | 0.04172 | 0.04362 | 10 | 3 |
| Reactome Prolonged ERK activation events | 0.04172 | 0.04362 | 10 | 3 |
| BioCarta Erk and PI-3 Kinase Are Necessary for Collagen Binding in Corneal Epithe | 0.04172 | 0.04362 | 10 | 3 |
| BioCarta Role of MEF2D in T-cell Apoptosis | 0.04172 | 0.04362 | 10 | 3 |
| GO:BP cell cycle checkpoint | 0.04393 | 0.04393 | 17 | 4 |

### Table S10. Enrichment tests for each type of ncRNA in different cell lines in Case Study 3

| | | | | | | |
|---|---|---|---|---|---|---|
| Fibroblasts | | lncRNAs | Other ncRNAs | | snRNAs | Other ncRNAs |
| | In network | 60 | 115 | In network | 41 | 134 |
| | Not in network | 789 | 893 | Not in network | 304 | 1378 |
| | | | **p-value=0.00187** | | | **p-value=0.103** |
| | | miRNAs | Other ncRNAs | | snoRNAs | Other ncRNAs |
| | In network | 17 | 158 | In network | 12 | 163 |
| | Not in network | 129 | 1553 | Not in network | 123 | 1559 |
| | | | p-value=0.419 | | | p-value=0.946 |
| HEK293T | | lncRNAs | Other ncRNAs | | miRNAs | Other ncRNAs |
| | In network | 51 | 92 | In network | 17 | 126 |
| | Not in network | 798 | 916 | Not in network | 129 | 1585 |
| | | | **p-value=0.0153** | | | p-value=0.0891 |
| | | snRNAs | Other ncRNAs | | snoRNAs | Other ncRNAs |
| | In network | 24 | 119 | In network | 16 | 127 |
| | Not in network | 321 | 1393 | Not in network | 119 | 1595 |
| | | | p-value=0.644 | | | **p-value=0.0871** |
| MCF7 | | lncRNAs | Other ncRNAs | | miRNAs | Other ncRNAs |
| | In network | 26 | 56 | In network | 8 | 74 |
| | Not in network | 21 | 216 | Not in network | 18 | 219 |
| | | | **p-value=1.232e-6** | | | p-value=0.702 |
| | | snRNAs | Other ncRNAs | | snoRNAs | Other ncRNAs |
| | In network | 14 | 68 | In network | 18 | 64 |
| | Not in network | 63 | 174 | Not in network | 47 | 190 |
| | | | p-value=0.113 | | | p-value=0.801 |

### Table S11. Computational time comparison between different methods (average over 100 replications)

| Scenario | Methods | | |
|---|---|---|---|
| High-dimension | | | |
| | | Mean (minutes) | Standard error (minutes) |
| | CAR-NET | 0.38 | 0.022 |
| | PC | 1.144 | 0.096 |
| | MMPC | 4.285 | 0.194 |
| | ARGES | 1.122 | 0.08 |
| | Pearson | 0.016 | 0.001 |
| Low-dimension | | | |
| | CAR-NET | 0.065 | 0.007 |
| | PC | 0.004 | 0.001 |
| | Hybrid MCM( | 0.591 | 0.003 |
| | GENIE3 | 2.236 | 0.003 |
| | HC | 0.007 | 0.001 |
| | MMPC | 0.008 | 0.001 |
| | ARGES | 0.004 | 0.001 |
| | Pearson | 0.001 | 0.001 |
